# βIII spectrin controls the planarity of Purkinje cell dendrites by modulating perpendicular axon-dendrite interaction

**DOI:** 10.1101/2020.02.14.945188

**Authors:** Kazuto Fujishima, Junko Kurisu, Midori Yamada, Mineko Kengaku

## Abstract

The mechanism underlying the geometrical patterning of axon and dendrite wiring remains elusive, despite its critical importance in the formation of functional neural circuits. Cerebellar Purkinje cell (PC) arborizes a typical planar dendrite, which forms an orthogonal network with granule cell (GC) axons. By using electrospun nanofiber substrates, we reproduce the perpendicular contacts between PC dendrites and GC axons in culture. In the model system, PC dendrites show preference to grow perpendicular to aligned GC axons, which presumably contribute to the planar dendrite arborization *in vivo*. We show that βIII spectrin, a causal gene for spinocerebellar ataxia type 5 (SCA5), is required for the biased growth of dendrites. βIII spectrin deficiency causes actin mislocalization and excessive microtubule invasion in dendritic protrusions, resulting in abnormally oriented branch formation. Furthermore, disease-associated mutations affect the ability of βIII spectrin to control dendrite orientation. These data indicate that βIII spectrin organizes the dendritic cytoskeleton and thereby regulates the oriented growth of dendrites with respect to the afferent axons.

**Summary statement:** βIII spectrin suppress the microtubule dynamics at the neuronal dendrite to inhibit the abnormal lateral branching, which causes misoriented branch formation.

## Introduction

Accurate information processing through the neuronal network requires proper development of dendrite pattern, which determines the receptive field, types of afferent fibers, and the number of synapses they form (London and Häusser, 2005). During development, neurons arborize their dendrites by a set of branch dynamics including extension, branching, and retraction, which are controlled by either cell-intrinsic programs or extrinsic cues from surrounding tissue (Dong et al., 2015; Jan and Jan, 2010; Valnegri et al., 2015).

Cerebellar Purkinje cells (PCs) develop highly branched dendrites in a single parasagittal plane. The dendrites are innervated by the parallel fiber axons of cerebellar granule cells (GCs), which run perpendicularly across the aligned PC dendrites along the coronal axis of the cerebellum. The planar dendrite with space-filling and no-overlapping arrangement is a distinguished feature of PCs and is thought to be advantageous for efficient network formation. For instance, planar dendrites connected with perpendicularly-oriented axonal bundles contribute to maximizing possible synaptic connections with minimal redundancy (Cuntz, 2012; Wen and Chklovskii, 2008). Indeed, each PC dendrite contacts with more than 100,000 parallel fibers in rodents (Napper and Harvey, 1988). PCs arborize their dendritic branches via dynamic remodeling during postnatal developmental stage (Fujishima et al., 2018; Joo et al., 2014; Kaneko et al., 2011; Kapfhammer, 2004; Sotelo and Dusart, 2009; Takeo et al., 2015; Tanaka, 2009). We have previously identified the basic rules of dendrite formation in PCs grown on coverslips (Fujishima et al., 2012). Typical fan-shaped branching structures of dendrites were constructed by constant extension and dichotomous terminal branching. Further, we have shown that retraction and stalling induced by dendro-dendritic contact (self-avoidance) play pivotal roles in the non-overlapping and space-filling distribution of dendritic branches (Fujishima et al., 2012; Kawabata Galbraith et al., 2018). Indeed, it has been shown that loss of cell-surface molecules such as clustered protocadherins and slit/robo, which recognize the contact between dendritic branches, affect the non-overlapping arrangement of PC dendrites (Gibson et al., 2014; Ing-Esteves et al., 2018; Kuwako and Okano, 2018; Lefebvre et al., 2012; Toyoda et al., 2014). Although these basic rules of dendritic growth can explain the dendritic configuration including the non-overlapping pattern on a 2D substrate, they cannot illustrate how dendrites acquire the planar arrangement in three-dimensional brain tissue.

Unlike other planar dendrites such as those of Drosophila DA neurons that extend along the structural boundary (Han et al., 2012; Kim et al., 2012), PC dendrites grow in a plane with no obvious scaffolds in the cerebellar cortex. Several lines of evidence suggest that the growth of PC dendrites in a direction perpendicular to the parallel fibers, which are formed earlier than the extension of PC dendrites (Altman and Anderson, 1972; Crepel et al., 1980; Nagata et al., 2006). The perpendicular growth of dendrite seems to be involved in the arborization of unique flat dendrites (Altman and Anderson, 1972; Crepel et al., 1980; Nagata et al., 2006; Sotelo and Dusart, 2009). Pioneering studies by Altman demonstrated that disruption of parallel fiber arrays via the X-ray irradiation of the developing cerebellum led to the disruption and realignment of PC dendrites in a direction perpendicular to the misoriented parallel fibers (Altman, 1973). However, the molecular machinery underlying the perpendicular growth of PC dendrites to the GC axon arrays is largely unknown (Gao et al., 2011; Kim et al., 2014).

Spectrins are structural molecules that form a tetrameric complex with two alpha and two beta subunits. Spectrin tetramers interact with actin filaments beneath the plasma membrane. Recent studies using super-resolution microscopy have revealed that spectrin, actin, and their associated proteins form membrane periodic skeletal (MPS) structures lining the circumference of the axons and dendrites (D’Este et al., 2015; Han et al., 2017; Vassilopoulos et al., 2019; Xu et al., 2013). In the MPS, ring-like structures of actin are connected by spectrin tetramers with a periodicity of ~190 nm. Alternatively, spectrin and associated proteins can form a 2D polygonal lattice structure in the soma or dendrites, possibly reminiscent of those found in erythrocytes (Han et al., 2017). These structures are considered to have multiple functions, such as the maintenance of membrane stiffness and/or elasticity (Krieg et al., 2017) and acting as a diffusion barrier to membrane molecules (Albrecht et al., 2016).

βIII spectrin, one of the beta subunits of spectrin molecules, is highly expressed in the PC dendrites and soma. βIII spectrin has been identified as a gene responsible for spinocerebellar ataxia type 5 (SCA5), which manifests as a progressive dysfunction of motor coordination (Ikeda et al., 2006). PCs in SCA5 patients and βIII spectrin knockout animals display aberrant dendrite morphologies due to abnormal arbor development and maintenance (Gao et al., 2011; Ikeda et al., 2006). Notably, the PCs of the βIII spectrin knockout mouse exhibit a defect in flat dendrite arborization (Gao et al., 2011), implicating βIII spectrin in the regulation of dendrite growth orientation. The loss of βIII spectrin leads to the mislocalization and/or diminished level of neurotransmitter receptors and transporters, which may cause the progressive neurodegeneration via excitotoxicity (Armbrust et al., 2014; Perkins et al., 2010; Stankewich et al., 2010). However, the mechanisms by which βIII spectrin controls dendrite development remain elusive.

Here, we show that βIII spectrin is indispensable for the directional arborization of PC dendrites perpendicular to parallel fibers, which is a prerequisite for the flat dendrite formation. We recapitulated a two-dimensional model of the orthogonal interaction between the dendrite and axonal bundles by using nanofibers as artificial scaffolds to reproduce parallel fiber arrays *in vitro.* We demonstrated that βIII spectrin-deficient PCs failed to form dendrites perpendicular to axonal bundles. Super-resolution observations confirmed the repeated lattice-like structures of βIII spectrin in PC dendrites. Loss of βIII spectrin caused abnormal cytoskeletal dynamics and misoriented growth of dendrites. Our data provide insights into the role of βIII spectrin in controlling the perpendicular connectivity of dendrites and axons to form planar dendrites.

## Results

### βIII spectrin is required for a cell-autonomous mechanism of planar dendrite arborization in PCs

Earlier studies have demonstrated that the planarity of PC dendrites is disturbed in loss-of-function mutants of βIII spectrin (Gao et al., 2011). We first confirmed the cell-autonomous role of βIII spectrin in PC dendrite formation by short hairpin RNA (shRNA) knockdown in wildtype background *in vivo*. We used a plasmid encoding an shRNA targeting βIII spectrin (βIII spectrin kd) that efficiently knocks down βIII spectrin expression in PCs in a dissociation culture (Fig. S1). We sparsely delivered the plasmid into PC precursors in the 4th ventricle via *in utero* electroporation at embryonic day 11.5 (E11.5). Consistent with previous observations using βIII spectrin knockout mice (Gao et al., 2011), we observed almost no differences in the total length of dendrites between control (ctr) and βIII spectrin knockdown cells at postnatal day 14 (P14) (Fig. 1A-C). Compared to the normal planar dendrites in control PCs, βIII spectrin knockdown cells exhibited misoriented dendrites that extended away from the main dendritic plane (Fig. 1A, B), similar to those in knockout animals (Gao et al., 2011). Indeed, quantitative analysis revealed that dendritic branches extruded from the main plane were significantly increased in βIII spectrin knockdown PCs (Fig. 1A, B, and D and Fig. S2, see Materials and Methods), suggesting that a cell-autonomous function of βIII spectrin is necessary for the planar dendrite formation of PCs.

**Fig. 1.**
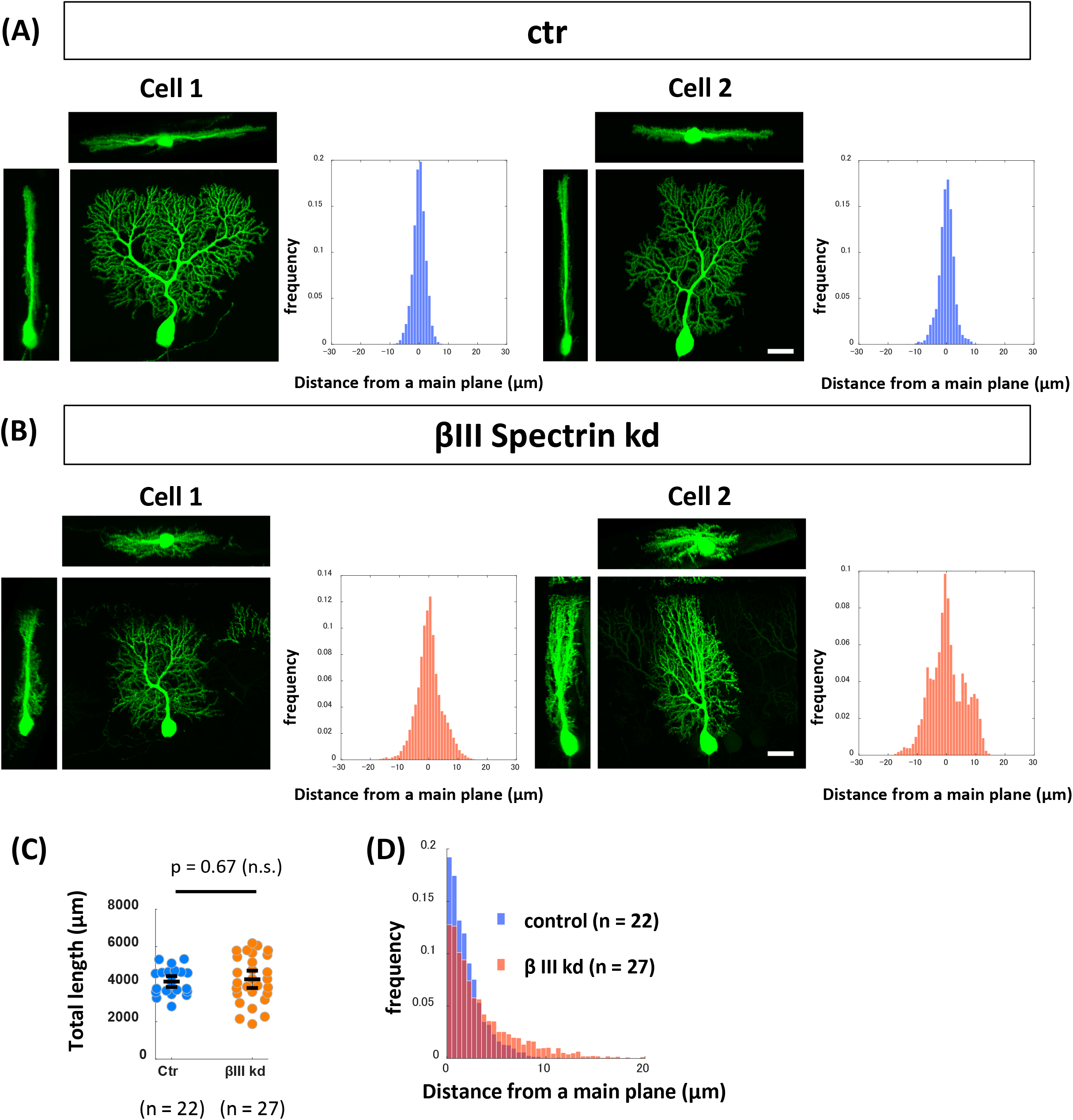
A Cell-autonomous role of βIII spectrin in the planar dendrite formation *in vivo*. **(A, B)** Morphology of PCs expressing the GFP/shRNA-control (A, ctr) or GFP/shRNA-βIII spectrin plasmid (B, βIII spectrin kd) in P14 sagittal sections. Top, front (sagittal) and side (coronal) views were taken from the three-dimensionally reconstructed images. Histograms showing the proportions of dendritic segments located at the indicated distances from the main dendritic plane. **(C)** Total dendritic length in control or βIII kd PCs at P14. Statistical analysis: Welch’s t-test. **(D)** Histograms showing the proportions of dendritic segments from the main dendritic plane. p = 2.7 x 10^-46^, two-sample Kolmogorov-Smirnov test. Scale bars: 20 μm

### Recapitulation of 2D perpendicular contact between PC dendrites and GC axons using artificial nanofibers *in vitro*

Next, we examined the mechanism underlying the effect of βIII spectrin deficiency on dendrite planarity. Considering that flat PC dendrites are closely aligned with neighboring PCs (Fig. 2A), it seems unlikely that gradients of guidance molecules instruct such dendritic configurations. It has previously been suggested that PC dendrites grow perpendicularly to GC axons (Nagata et al., 2006). If so, dendrites will extend in parasagittal planes perpendicular to the bundle of parallel fibers (granule cell axons) that run along the coronal axis (Fig. 2B, left). We thus speculated that βIII spectrin might be involved in dendrite growth perpendicular to GC axons.

**Fig. 2.**
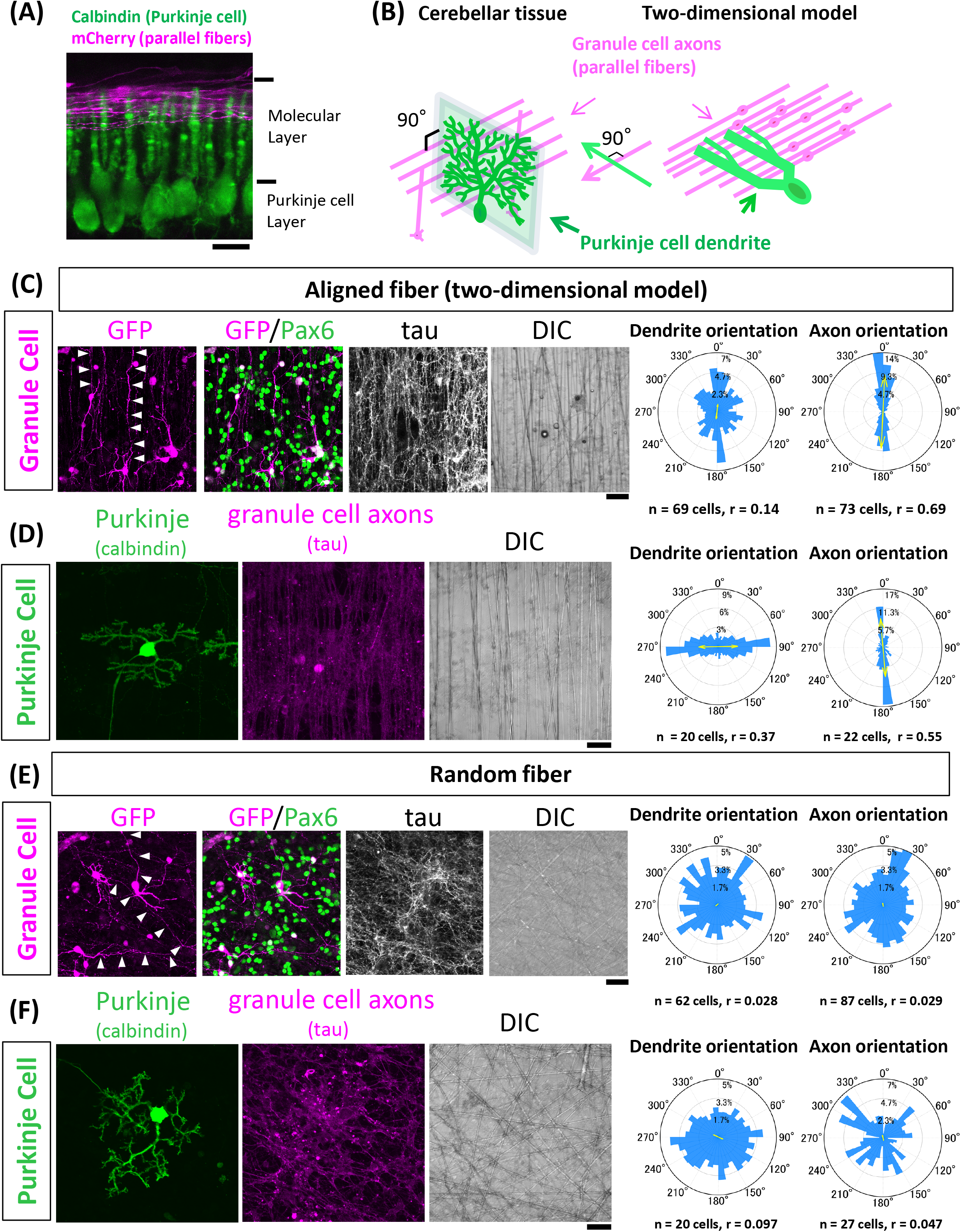
Reconstruction of perpendicular interactions between PC dendrites and GC axons using artificial nanofibers. **(A)** PCs (calbindin) and parallel fibers (mCherry, introduced by *in vivo* electroporation) in the P11 coronal section. **(B)** Schematic illustrations showing the arrangement of PC dendrites and parallel fibers (GC axons) in cerebellar tissue and a corresponding 2D model. **(C-F)** Representative images of GCs (C, E) and PCs (D, F) grown on nanofibers (C, D: aligned and E, F: random). In (C) and (E), Pax6-positive GC morphologies were visualized with GFP. Axonal bundles were stained with tau. In (D) and (F), PCs were visualized by calbindin staining. Axonal bundles were stained with tau. The polar histograms indicate the angular distribution of the axons or dendrites of GCs or PCs. The mean vector is shown with the yellow arrow, and its length is indicated as r. Scale bars: 30 μm

To test this hypothesis, we first confirmed whether normal PC dendrites preferentially grow perpendicularly to GC axons. We established a simplified model in which parallel GC axons are recapitulated in 2D spaces (Fig. 2B, right). We utilized electrospun polycaprolactone nanofibers, which have been used to navigate axonal extensions in culture (Hyysalo et al., 2017). We plated dissociated cells from P0 cerebella on aligned or randomly oriented nanofibers. As expected, GC axons labeled with GFP ran parallel to each other on aligned nanofibers (arrowheads in Fig. 2C), while they extended randomly on non-oriented nanofibers (arrowheads in Fig. 2E).

We previously reported that PCs in dissociated cultures initiate dendrites at approximately 8 days *in vitro* (8 DIV) and radially extend branches until at least 14 DIV (Fujishima et al., 2012). Likewise, the PC dendrites on the randomly oriented nanofibers were observed to grow radially with no directional preference (Fig. 2F). In contrast, the distribution of PC dendrites on the aligned nanofibers was highly biased toward the direction perpendicular to fibers, supporting the notion that PC dendrites grow perpendicularly to GC axons (Fig. 2D). Indeed, quantitative analysis revealed that the majority of dendritic segments oriented perpendicular to aligned fibers (polar histogram: dendrite orientation in Fig. 2D, see Materials and Methods, and Fig. S3). The perpendicular growth was specific to PC dendrites, as PC axons grew parallel to GC axons (Fig. 2D). Furthermore, GC dendrites grew in a random orientation on aligned fibers (Fig. 2C). These results suggest that the perpendicular interaction is not ubiquitous but rather specific to certain synaptic partners, including PC dendrites and GC axons.

To verify that PC dendrites are navigated by GC axons but not by nanofibers, we confirmed the interactions between PC dendrites and GC axons without nanofibers (Fig. S4A). We prepared microexplants of the external granule cell layer populated with differentiating GCs from the P2 cerebellum (Kawaji et al., 2004; Nagata and Nakatsuji, 1990). Isolated PCs were cocultured at 1 DIV on the GC axons that extended radially from the explant. We confirmed that PCs developed perpendicular dendrites to the radially extended GC axons at 10 DIV (Fig. S4A), consistent with a previous report (Nagata et al., 2006). Surface rendering showed close apposition of the dendrites and axons (Fig. S4B). These results suggest that direct PC dendrite-GC axon interaction determines the dendrite orientation.

To further exclude the possibility that the nanofibers act as scaffolds for PC dendrite growth, we prepared PC cultures with GCs of different densities on nanofibers (Fig. S5). Although PCs established perpendicular dendrites in high- and medium-density cultures, they could not grow dendrites on free nanofibers devoid of GC axons in a low-density culture, suggesting that nanofibers do not serve as growth scaffolds (Fig. S5A, B). Taken together, we conclude that PC dendrites grow perpendicular to GC axons.

### Knockdown of βIII spectrin interferes with perpendicular contact

To examine whether βIII spectrin is required for the perpendicular contact between PC dendrites and GC axons, we introduced shRNA plasmids to PCs and plated them on nanofiber substrates. In contrast to the dendrites perpendicular to GC axons observed in control cells, a significant proportion of the dendritic branches in βIII spectrin knockdown cells extended in different orientations (Fig. 3A). Quantitative morphometry revealed that the total length and branch number were increased in βIII spectrin knockdown cells, suggesting the excess formation of misoriented dendrites (Fig. 3B, C). Coexpression of shRNA-resistant βIII spectrin reversed the changes in dendrite orientation, total length, and branch number, negating the off-target effects (Fig. 3A-C, βIII kd + rescue). Furthermore, the CRISPR/Cas9-based βIII spectrin knockout PCs exhibited similar phenotypes (Fig. S6). These results indicate that βIII spectrin is required for perpendicular contact between PC dendrites and GC axons.

**Fig. 3.**
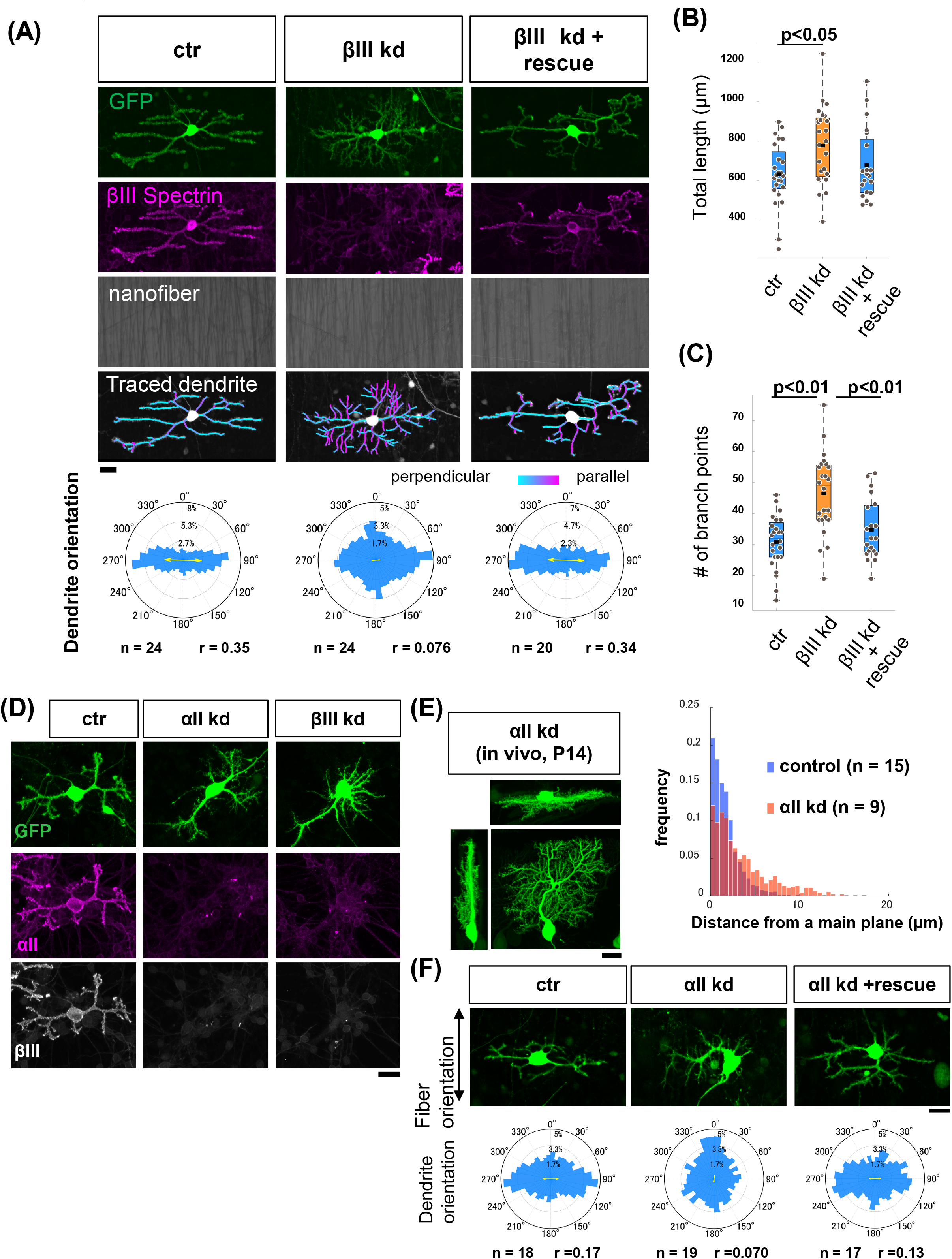
Disruption of perpendicular dendrite growth via the knockdown of βIII spectrin. **(A)** Morphology of PCs transfected with GFP/shRNA control (left panels, ctr), GFP/shRNA-βIII spectrin (middle panels, βIII kd) and GFP/shRNA-βIII spectrin together with shRNA-resistant βIII spectrin (right panels, βIII kd + rescue) grown on aligned fibers (12 DIV). βIII spectrin signals were shown in magenta. Traced dendrites are pseudocolored based on the angles between the dendritic segments and fibers. The polar histograms indicate angular distributions of dendritic segments. p = 6 x 10^-16^ for ctr vs βIII kd, p > 0.5 for ctr vs rescue, and p = 6 x 10^-16^ for βIII kd vs rescue (Watson-Wheeler test with Bonferroni correction). **(B, C)** Quantification of total dendritic length (B) and the total branch number (C) in control or βIII kd PCs. Statistical analysis: ANOVA followed by the Tukey-Kramer post hoc test. **(D)** Morphology of PCs transfected with GFP/shRNA-control (ctr), GFP/shRNA-αII spectrin (αII kd) and GFP/shRNA-βIII (βIII kd). αII and βIII spectrin signals were shown in magenta and white, respectively. **(E)** Morphology of PCs transfected with GFP/shRNA-αII spectrin (αII kd) in P14 sagittal section. The histogram shows the proportions of dendritic segments located at the indicated distance from the main dendritic plane. p = 2.2 x 10^-39^, two-sample Kolmogorov-Smirnov test. **(F)** Top panels: Morphology of PCs transfected with GFP/shRNA-control (ctr), GFP/shRNA-αII spectrin (αII kd), and GFP/shRNA-αII spectrin plus shRNA-resistant αII spectrin (αII kd + rescue) grown on aligned nanofibers. Bottom panels: Angular distribution of dendritic segments. Scale bars: 20 μm.

Spectrin molecules form tetrameric complexes composed of α and β subunits, and the loss of α subunits destabilizes β subunits in embryonic tissue (Stankewich et al., 2011). Among α-spectrin subtypes, αII spectrin is most abundant in non-erythrocytic cells (Cianci et al., 1999; Winkelmann and Forget, 1993). Indeed, αII spectrin was strongly expressed in the PC somata and dendrites, similar to βIII spectrin (Fig. 3D). We found that the knockdown of either αII or βIII spectrin dramatically reduced both proteins in PCs, suggesting the interdependency of the molecules in the stabilization of the spectrin complex (Fig. 3D). Accordingly, αII spectrin knockdown disrupted planar arborization *in vivo* (Fig. 3E) as well as biased dendrite organization on aligned nanofibers (Fig. 3F). Thus, these results indicate that the αII/βIII spectrin complex regulates dendrite orientation in PCs.

### Loss of βIII Spectrin causes aberrant branch formation

We have previously demonstrated that the non-overlapping arrangement of PC dendrites is achieved by the contact-dependent retraction of growing branches (Fujishima et al., 2012). Thus, the perpendicular dendrites on nanofibers might be attributed to either (1) biased outgrowth (extension and branch formation) in the perpendicular direction or (2) the biased retraction of misoriented branches. To distinguish between these possibilities, we performed time-lapse observations of growing dendrites on aligned nanofibers. We traced the outgrowth of PC dendrites (~0.7 μm/hour) every three hours for several days from 8 DIV when dendrite formation was initiated in the culture (Fujishima et al., 2012). Control PCs continuously grew their dendrites by extension and branch formation perpendicular to the GC axon orientation (Fig. 4A, B). Next, we measured the orientation of retracted branches. Dendritic retractions were triggered by repetitive collisions of growing dendritic tips in agreement with our previous report (Fujishima et al., 2012). Of all dendritic retractions observed in control PCs, about 86% of them were induced after obvious dendritic collisions. In addition, about 73% of branches were eliminated after a collision in control PCs (Fig. 4D). Disorienting branches were eliminated with high probability as they had more chance to contact with other branches, yet perpendicular branches were also retracted when they collided with other growing branches (Fig. 4C). Therefore, we concluded that the perpendicular dendrite orientation in control PCs was likely due to biased dendritic growth, although the elimination of disoriented branches may contribute to some extent.

**Fig. 4.**
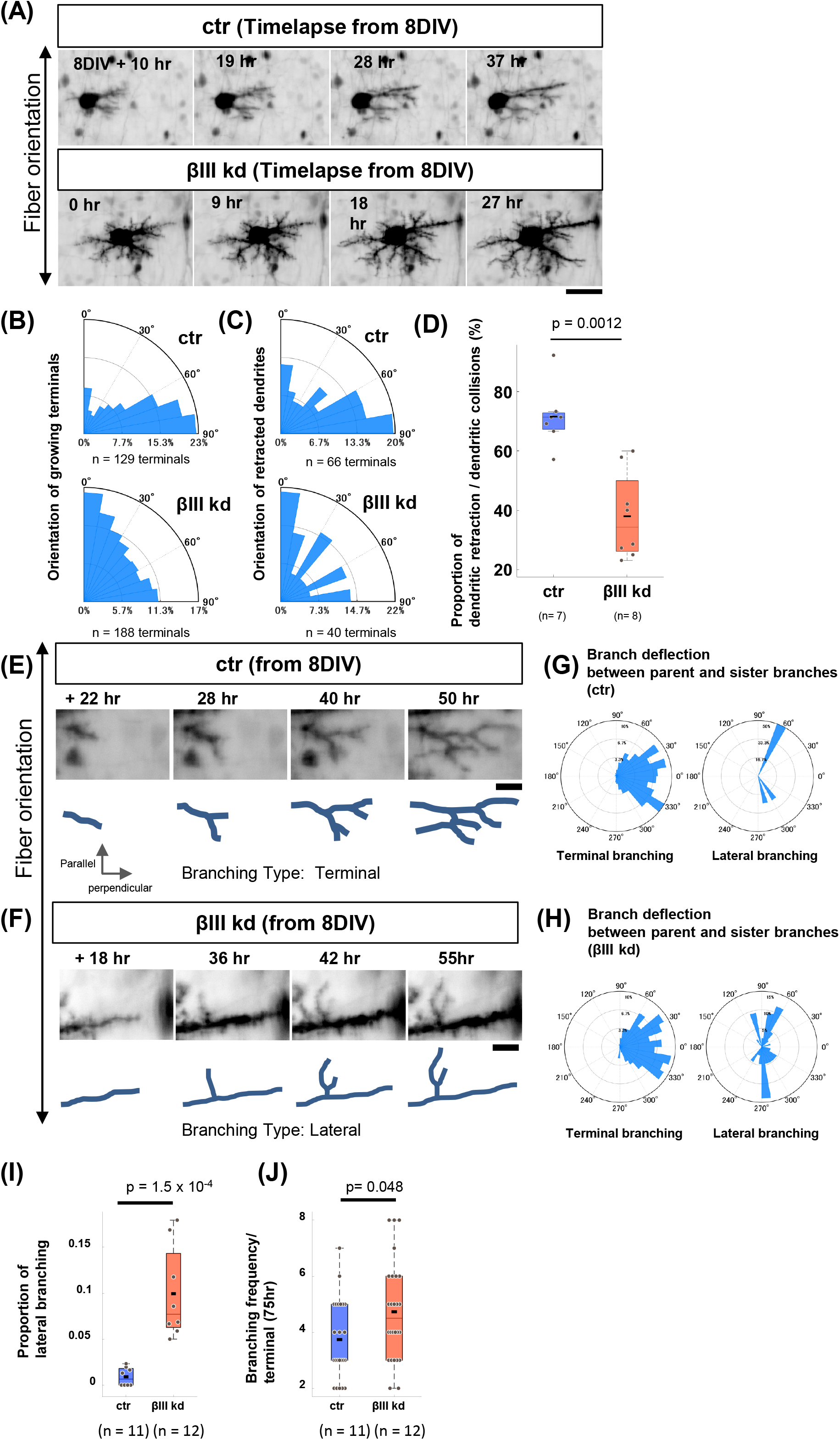
Analysis of PC dendrite dynamics on aligned nanofiber substrates. **(A)** Time-lapse images of developing dendrites in GFP/shRNA-control (ctr) or GFP/shRNA-βIII spectrin (βIII kd) PCs growing on aligned nanofibers from 8 DIV. **(B, C)** Polar histograms showing the angular distribution of growing dendritic terminals (B) or retracted dendrites (C). **(D)** The number of retracted dendrites per cell during the 75-hr time-lapse observation. Statistical significance: Wilcoxon rank-sum test. **(E, F)** Magnified views of time-lapse images of control (E) or βIII kd (F) dendrites focusing on the branch formation. The images show the examples of terminal branching (E) or lateral branching (F) observed in control or βIII kd cells, respectively. Branch formation of more than 15 μm away from the terminal was defined as lateral branching. **(G, H)** Polar histograms showing the deflection angle between parent and sister branches at terminal branching (left) or lateral branching (right) in ctr (G) and βIII kd cells (H). n = 366 branching events for ctr and n = 429 branching events for βIII kd. **(I, J)** Box graphs showing the proportion of lateral branching (I) and the branching frequency of growing dendrites (J). Statistical analysis: Wilcoxon rank-sum test. Scale bars: 40 μm in (A), 10 μm in (C) and (D).

In βIII spectrin knockdown cells, the fraction of parallel-growing dendritic terminals was markedly increased (Fig. 4A, B). Similar to control PCs, most of the dendritic retractions (~90%) in βIII spectrin knockdown cells occurred after the dendritic collisions. However, the frequency of dendrite retraction after a collision was reduced to 39% in βIII spectrin knockdown cells, suggesting that the contact-dependent dendritic retraction is dysregulated in βIII spectrin deficient cells (Fig. 4D). However, the orientation of retracted dendrites was strongly biased parallel to the axons, negating that the increase in misoriented dendrites in βIII spectrin knockdown cells was caused by suppression of dendrite retraction of wrong arbors. Therefore, we focused on how βIII Spectrin deficiency leads to the dysregulation of dendrite growth orientation.

Our previous studies have shown that PCs form dendritic branches primarily via the bifurcation of growing terminals (terminal branching), while they rarely extend collaterals from the shaft (lateral branching) (Fujishima et al., 2012). Accordingly, control PC dendrites on aligned fibers mainly displayed terminal branching (Fig. 4E, I). In contrast, lateral branching was increased by more than 10-fold in βIII spectrin knockdown dendrites compared with control dendrites (Fig. 4F, I), although the branching frequency was only slightly altered (Fig. 4J). The deflection angle between the bifurcated terminal branches ranged within approximately ±30°, while that of lateral branching was greater than 60° in both the control and βIII spectrin deficient dendrites (Fig. 4G, H). These results suggest that βIII spectrin regulates the perpendicular growth of dendrites by inhibiting lateral branching. Thus, the frequent lateral branching may contribute to the increase in misoriented dendrites in the βIII spectrin deficient cells.

### Lateral branch formation in βIII spectrin-knockdown dendrites

We next observed how dendritic planarity was affected in βIII spectrin knockdown cells *in vivo*. Control PC dendrites transfected with GFP aligned in a parasagittal plane in the molecular layer parallel to the neighboring PC dendrites (Fig. 5A, B). These PCs rarely exhibited dendritic branches growing in lateral (coronal) directions. Neighboring dendrites were separated by gaps of approximately 1-3 μm, showing minimal crossing with adjacent branches. Similar to control PCs, the main dendritic arbors of βIII spectrin knockdown PCs were mostly parallel to neighboring PC dendrites in coronal sections (Fig. 5A, B, βIII kd), although some dendrites bent or tilted into incorrect planes (white arrows in Fig. 5A). Notably, βIII spectrin knockdown dendrites exhibited an increased number of laterally oriented branches growing into the territories of the neighboring PC dendrites (yellow arrows in Fig. 5A, Fig. 5C). These misoriented branches often turned and extended parasagittally into the gaps between PC dendrites (yellow arrowhead in Fig. 5A). These misoriented lateral branches likely contribute to the disruption of planar dendrite in βIII spectrin deficient PCs.

**Fig. 5.**
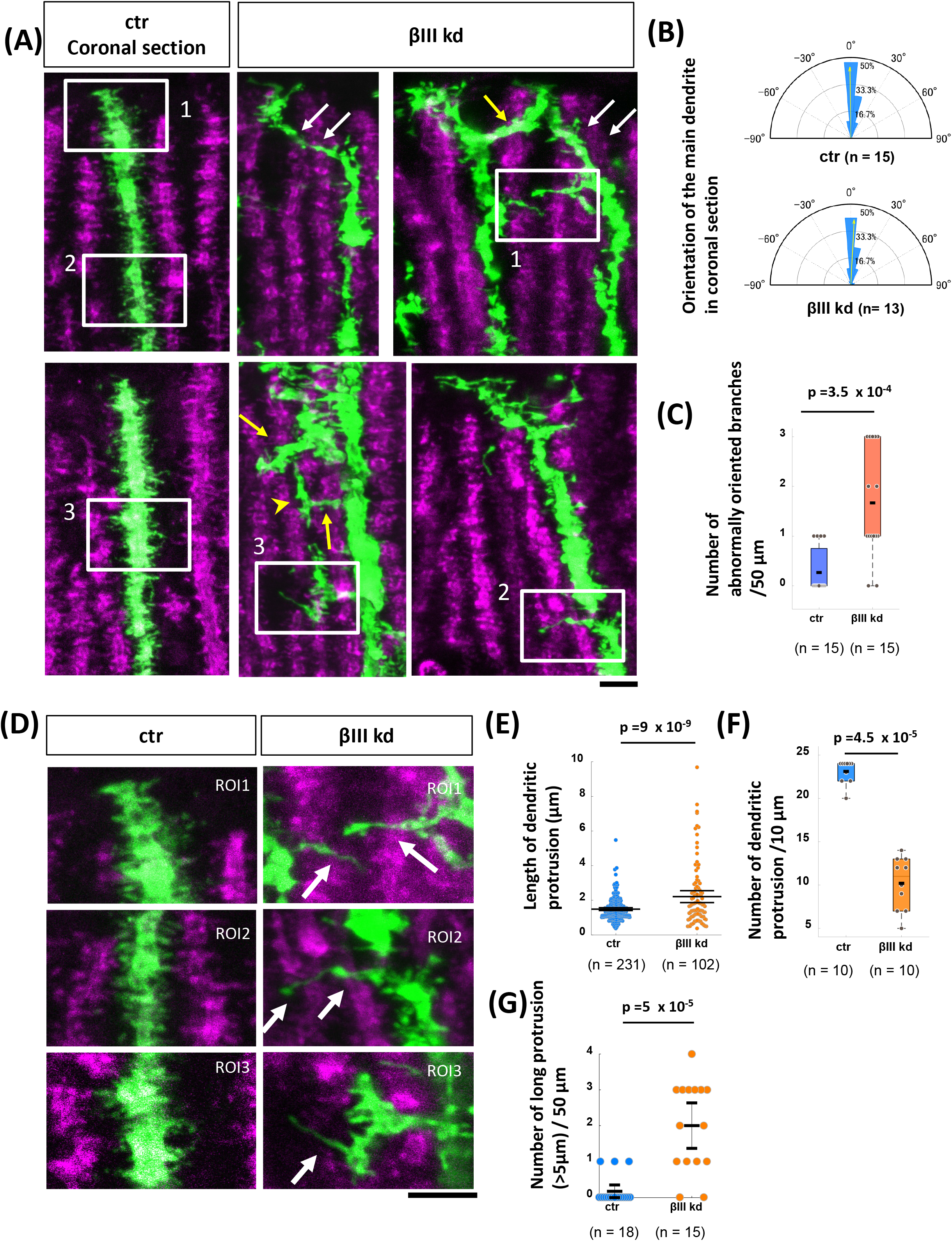
βIII spectrin knockdown induces the extension of lateral-oriented branches. **(A)** Growing dendritic arbors in GFP/shRNA-control (ctr) or GFP/shRNA-βIII spectrin (βIII kd) transfected PCs in the vermis region in P14 coronal slices. βIII spectrin signals were shown in magenta. **(B)** Polar histogram showing the angular distribution of the main dendritic arbor in control and βIII kd cells. **(C)** Quantification of the number of laterally oriented branches extruded from main dendrites. **(D)** Images are magnified views of the insets in (A). **(E)** Quantification of the length of dendritic protrusions in control or βIII kd cells. **(F)** Number of dendritic protrusions per 10 μm of the dendritic segment at the distal region in control or βIII kd cells. **(G)** The number of abnormally long (> 5 μm) dendritic protrusions per 50 μm of the dendritic segment in control or βIII kd cells. Statistical analyses: Wilcoxon rank sum test (C) and Student’s t-test (E-G). Scale bars: 5 μm in (A) and (D).

It has been demonstrated that the growing PC dendrites are covered with numerous dendritic protrusions, including dendritic filopodia and immature spines (Kawabata Galbraith et al., 2018; Shimada et al., 1998). As dendritic protrusions are known to serve as dendritic branch precursors in some neurons, we next analyzed the dendritic protrusions in PCs with or without βIII spectrin expression. Control dendrites presented numerous dendritic protrusions emanating from the shaft with a mean length of 1.48 ± 0.04 μm (mean ± SEM, n = 231) (Fig. 5D, E). In contrast, βIII spectrin knockdown dendrites exhibited significantly longer protrusions (2.21 ±0.17 μm, n = 102) at a lower density, in agreement with previous studies (Fig. 5D-F) (Efimova et al., 2017; Gao et al., 2011). Notably, some dendritic protrusions in βIII spectrin knockdown cells were abnormally elongated (>5 μm) in lateral directions away from the main sagittal plane of the dendritic shaft (arrows in Fig. 5D, Fig. 5G). These long lateral protrusions seemed to serve as precursors of the ectopic lateral branches in βIII spectrin deficient cells.

### Abnormal formation of dendritic protrusions in βIII spectrin-knockdown dendrites

To further analyze the implications of βIII spectrin in the formation of dendritic protrusions and branches, we observed the dendritic structures in neurons grown on aligned nanofibers. Control PCs bore highly dense protrusions that covered the lateral surface of the dendritic shaft, similar to those observed *in vivo* (Fig. 6A). These protrusions expressed glutamate receptor δ2 (GluD2), which functions as a synaptic glue by binding with presynaptic neurexin and cbln1 in GC axonal terminals (Matsuda et al., 2010; Uemura et al., 2010), suggesting that these protrusions are dendritic spines or immature spine precursors (Fig. 6A).

**Fig. 6.**
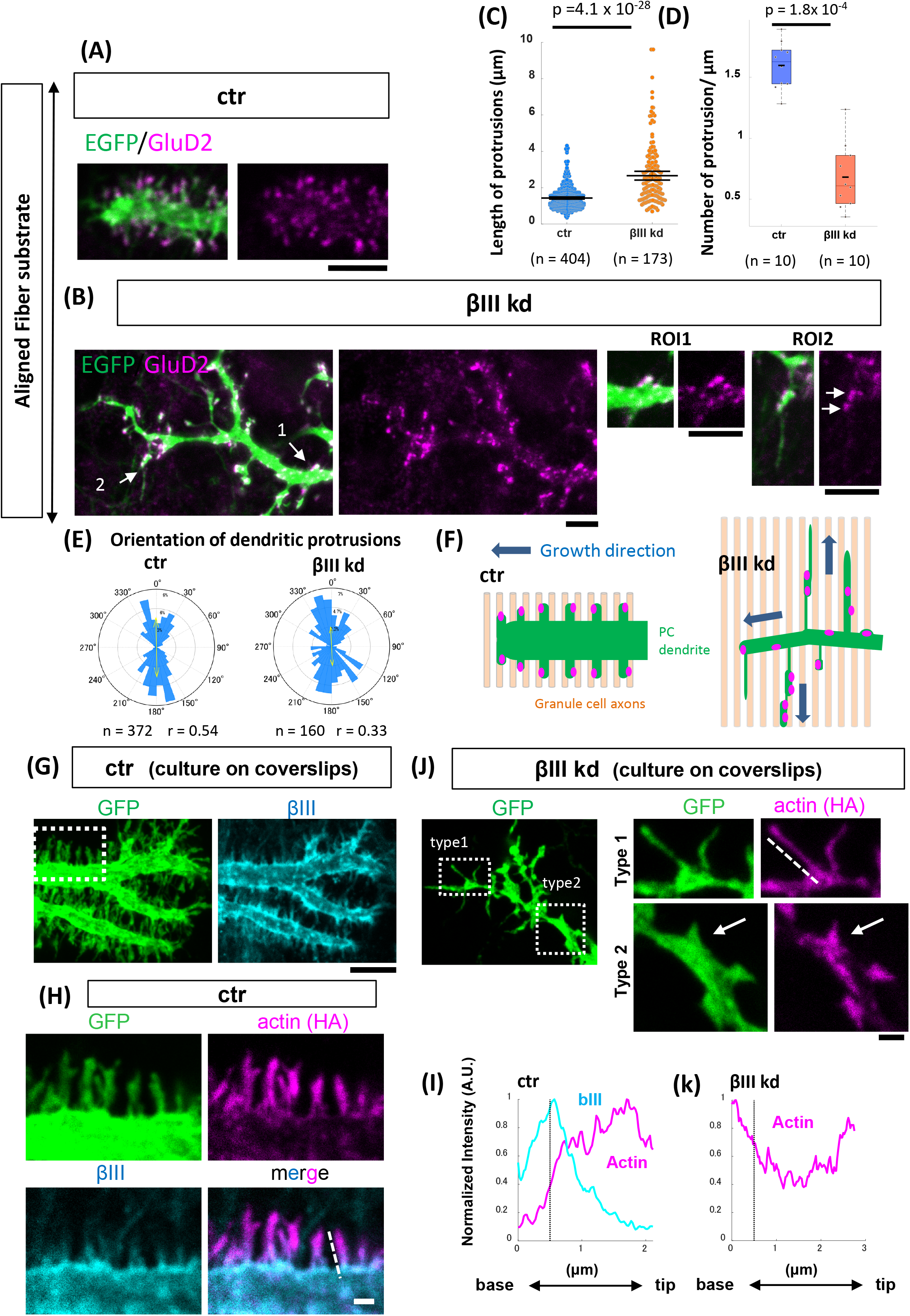
Abnormal extension of dendritic protrusions is induced by the loss of βIII spectrin. **(A, B)** Representative images of the growing dendritic terminals of PCs transfected with GFP/shRNA-control (A, ctr) or GFP/shRNA-βIII spectrin (B, βIII kd). Cells were cultured on aligned nanofibers and stained with the postsynaptic protein GluD2 (magenta) at 10 DIV. ROI1 and RO12 were magnified views of the regions indicated by arrow1 and 2 in the left image. **(C, D)** Quantification of the length of the dendritic protrusions (C) and the number of dendritic protrusions per micron (D) in control or βIII kd PC dendrites. Statistical analyses: Wilcoxon rank-sum tests. **(E)** The orientation of dendritic protrusions in ctr or βIII kd PC dendrites. **(F)** Schematic explanation of growing control or βIII kd PC dendrites (green) on aligned GC axons (light orange). **(G)** Localization of βIII spectrin (cyan) at the distal end of the growing dendrite in a control PC grown on a coverslip. **(H)** Magnified views of actin and βIII spectrin signals at the dendritic filopodia in the boxed region of (G). HA-actin was transfected to visualize actin signals. **(I)** Intensity profiles of βIII spectrin and HA-actin along the dotted line marking a dendritic protrusion in (H). The black dotted line in the graph indicates the boundary between the protrusion and the shaft. **(J)** Left: Representative image of the distal end in the growing dendrite of a GFP/βIII spectrin kd PC grown on a glass substrate. Right: Magnified views of actin signals in the dendritic protrusions (Type 1: thin and long, Type 2: short and stubby) of βIII kd cells. **(K)** Intensity profiles of HA-actin along the dotted line marking the growing terminal of the dendrites in (J). Scale bars: 5 μm in (A), (B), (G) and (J, left), 1 μm in (H) and (J, right panels).

Compared to control cells, the distal dendrites of βIII spectrin knockdown cells were significantly thinner (ctr: 1.80 ± 0.15 μm, n= 19, βIII kd: 0.90± 0.06 μm, n= 22, mean ± SEM, p = 7 x 10^-6^, Student’s t-test) (Fig. 6B). In contrast to the dendritic protrusions in control cells, which presented a relatively constant length of 1-3 μm, those in βIII spectrin knockdown cells presented various lengths at a lower density, with an average length that was significantly longer than that in control cells (Fig. 6C, D). We found some abnormally elongated protrusions of nearly 10 μm in βIII spectrin knockdown dendrites that extended parallel to the orientation of GC axons. Other short protrusions often appeared wider at their bases, reminiscent of the shaft synapses with deficient neck formation observed in βIII spectrin deficient hippocampal neurons (Fig. 6B ROI1)(Efimova et al., 2017). The extremely long protrusions exhibited multiple GluD2 puncta that were irregularly arranged along their lengths (Fig. 6B ROI2). The orientation of the protrusions in the control and βIII spectrin knockdown dendrites was mostly parallel to GC axons (Fig. 6E). These results suggested that dendritic protrusions in βIII spectrin deficient dendrites abnormally extend along GC axons, which possibly induce disoriented branch formation (Fig. 6F).

We next analyzed the subcellular localization of βIII spectrin in growing dendrites in PCs cultured on coverslips. In agreement with previous reports (Efimova et al., 2017; Gao et al., 2011), βIII spectrin was strongly localized on the surface of the dendritic shaft and the base of dendritic protrusions, while it was excluded from the protrusion tips (Fig. 6G, H). Actin showed an inverse gradient along the dendritic protrusions such that it was densely localized at the tip and sharply declined in the base of the protrusion and the dendritic shaft (Fig. 6H, I). In contrast, in βIII spectrin knockdown PCs, actin was more widely distributed along the entire length of both long and thin (Type 1 in Fig. 6J) and short and stubby (Type 2 in Fig. 6J) dendritic protrusions and was often dispersed in the shaft of distal dendrites (Type 1 in Fig. 6J, Fig. 6K). These results imply that βIII spectrin might be involved in the formation of the structural boundary between the dendritic shaft and protrusions that confines actin filaments within dendritic protrusions.

### Membrane periodic skeleton structure formed by βIII spectrin

Super-resolution microscopy has revealed that the spectrin/actin complex forms membrane periodic skeletal (MPS) structures, which might function as a diffusion barrier for membrane proteins (Albrecht et al., 2016; Leite et al., 2016). Using stimulated emission depletion microscopy (STED), we confirmed the existence of MPS-like repeated structures composed of βIII spectrin in the shaft regions of developing PC dendrites (Fig. S7A, C: ROI1). The average interval of the repeated structures was 186 ± 5 μm (Fig. S7D), consistent with previous studies (Xu et al., 2013). Actin rings were not clearly observed in the repeated structures, probably due to very low actin signals in the dendritic shafts compared to dendritic filopodia (data not shown and Fig. 6H). Repeated structures of βIII spectrin were also observed along the narrow corridor of the neck region of dendritic protrusions (Fig. S7B, C: ROI2 and 3) with an interval of 187 ± 5 μm (Fig.S7D). The repeated spectrin structures were not continuous but were often interrupted by irregularly arranged subsets, consistent with previous observation demonstrating the lower propensity for repeated structure formation in dendrites (Fig. S7E, F)(D’Este et al., 2015; Han et al., 2017). Thus, βIII spectrin formed random meshwork or repeated structures in dendritic shafts and filopodial bases.

### βIII spectrin suppresses microtubule entry into dendritic protrusions

It has previously been demonstrated using young hippocampal neurons that the transition from filopodia to neurites is triggered by microtubule invasion into the filopodia following local actin remodeling (Dent et al., 2007; Flynn et al., 2012). To test whether βIII spectrin contributes to blocking microtubule entry into dendritic protrusions, we monitored microtubule polymerization in growing dendrites by transfecting EB3-EGFP, a plus-end marker of dynamic microtubules. We analyzed the growing terminals of dendrites (within 10 μm from the terminal) and more proximal dendrites (more than 10 μm away from the terminal) separately to examine the regional difference in EB3 dynamics.

In control cells, 48 ± 5% (mean ± SEM, 226 protrusions from 18 dendrites) of dendritic protrusions around the growing terminal were targeted by EB3-EGFP within 150 seconds of observation (Fig. 7A, C). In contrast, a significantly lower proportion of protrusions were invaded by EB3 in proximal dendrites (5 ± 2%, 149 protrusions from 11 dendrites), in line with the notion that microtubule entry into filopodia triggers neurite extension at dendritic tips.

**Fig. 7.**
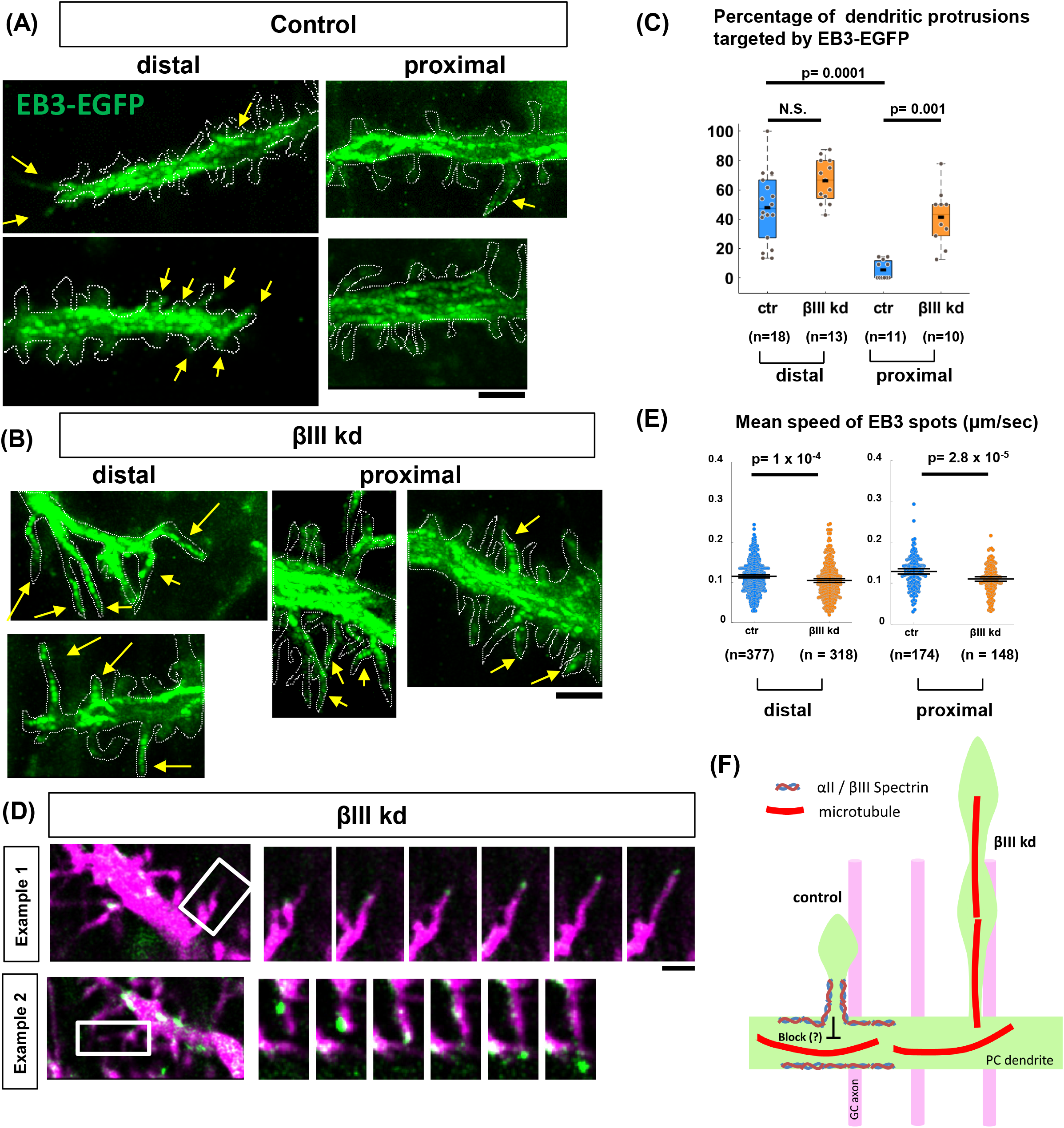
The invasion of EB3 into dendritic protrusions is facilitated by the loss of βIII spectrin. **(A, B)** Time-lapse imaging of EB3-EGFP in the dendrites of mCherry/shRNA-control (A, ctr) or mCherry/shRNA-βIII spectrin (B, βIII kd) PCs. Images are the maximum projections of EB3-EGFP spots during 150 sec of observation. The most distal (left) and the proximal (right) dendritic regions were recorded every three seconds. The dotted line indicates the contour of the dendrites (mCherry). EB3-EGFP signals in the dendritic protrusions are indicated by arrows. **(C)** The ratio of filopodia targeted by EB3-EGFP within 150 sec of observation. Statistical analysis: Steel-Dwass multiple comparison test. **(D)** Example of filopodial extension triggered by EB3-labeled microtubule polymerization in the proximal dendrite region in βIII kd cells. Right panels show time-series images (12-sec intervals) of the boxed region in the left images. **(E)** Quantification of the mean speed of EB3 spots traveling within PC dendrites. Statistical analysis: Wilcoxon rank-sum test. **(F)** The schematic hypothesis of βIII spectrin function in the regulation of microtubule dynamics. Scale bars: 3 μm in (A) and (B), 2 μm in (D)

We found that βIII spectrin knockdown significantly increased the proportion of EB3-targeted protrusions in proximal regions (41 ± 6%, 91 protrusions from 10 dendrites) (Fig. 7B, C), while only a slight increase was observed in the distal area (67 ± 4%, 102 protrusions from 13 dendrites). These data support the idea that βIII spectrin interferes with microtubule invasion into dendritic protrusions in proximal dendrites. Furthermore, we often observed that EB3-positive puncta tipped the filopodia and promoted their aberrant extension in βIII spectrin knockdown cells (Fig. 7D). The speed of microtubule polymerization was instead slightly downregulated in βIII spectrin-knockdown cells, negating that excessive microtubule entry in βIII spectrin-deficient cells was caused by increased microtubule polymerization activity (Fig. 7E). These results suggest that βIII spectrin controls directed dendritic arborization by suppressing microtubule invasion and ectopic branch formation from proximal dendritic protrusions (Fig. 7F).

### Mutations causing spinocerebellar ataxia type 5 (SCA5)

Mutations in βIII spectrin are known to cause spinocerebellar ataxia type 5 (SCA5). Hence, we wondered if the disease mutations affect biased dendrite growth perpendicular to GC axons. We focused on three mutations identified in earlier studies (Fig. 8A) (Ikeda et al., 2006). The first is a point mutation found in a German family that results in a leucine to proline substitution (L253P) in the calponin homology domain; the second is an in-frame 39-bp deletion (E532-M544 del) in the third spectrin repeat found in an American family; and the third is a 15-bp deletion in the third spectrin repeat (5-amino acid deletion with the insertion of tryptophan, L629-R634 delinsW) found in a French family (Fig. 8A). The amino acid sequences related to these mutations are conserved among humans and mice. Thus, we generated mouse βIII spectrin mutant constructs harboring the corresponding mutations. These constructs were designed to be shRNA resistant and were tagged with a myc epitope at their N-termini for imaging.

**Fig. 8.**
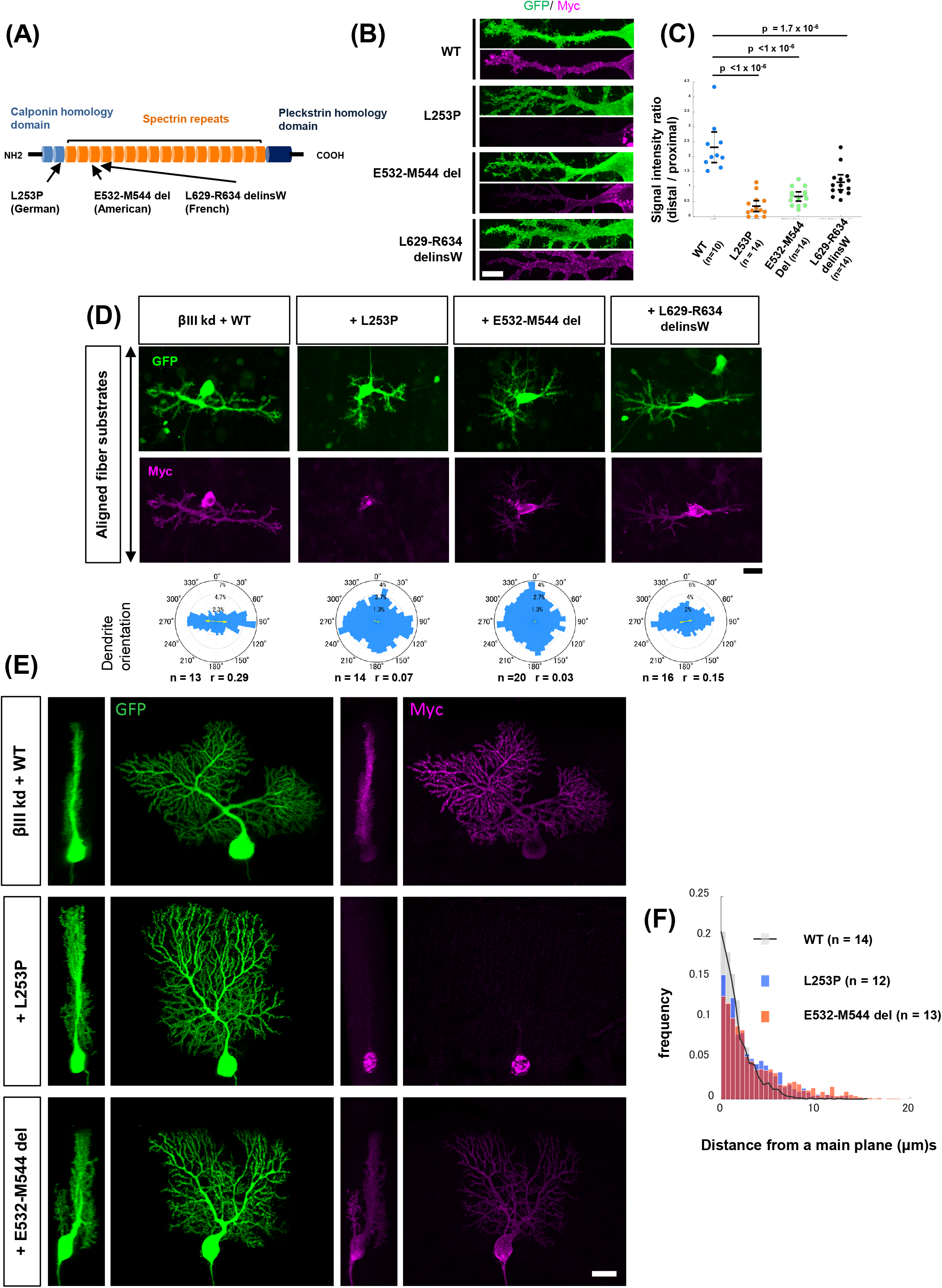
Functional analysis of the effect of SCA5-related mutations on the regulation of dendrite orientation. **(A)** Domain structure of βIII spectrin and familial mutations associated with SCA5. **(B)** Localization of myc-tagged βIII spectrin wild-type (WT), L253P, E532-M544 del, and L629-R643 delinsW mutant proteins in PCs grown on coverslips (9 DIV). **(C)** Quantification of the signal intensity ratio (distal versus proximal) of βIII molecules in PC dendrites. Statistical analysis: Kruskal-Wallis test followed by the Dunnett test. **(D)** Morphologies of PCs transfected with GFP/shRNA-βIII spectrin plus myc-tagged βIII spectrin wild-type (WT), L253P, E532-M544 del, and L629-R643 delinsW mutant proteins grown on aligned nanofibers. Polar histograms indicate the angular distribution of dendritic segments. (E) Morphologies of PCs expressing the GFP/shRNA-βIII plasmid with myc-tagged βIII spectrin wildtype (WT), L253P, or E532-M544 del mutant in P14 sagittal slices. (F) Histograms showing the proportions of dendritic segments located at the indicated distances from the main plane. Scale bars: 10 μm in (B), 20 μm in (D) and (E).

Dissociated PCs on aligned nanofiber substrates were knocked down for endogenous βIII spectrin and concomitantly transfected with wild-type βIII spectrin or disease-related mutants. Wild-type βIII spectrin was distributed in the somatodendritic area up to the most distal dendritic regions. In sharp contrast, the L253P mutant form was localized in intracellular vesicular structures in the somatic area, while almost no signal was observed in dendrites (Fig. 8B). On the other hand, mutants with deletions in the third spectrin repeat (E532-M544 del and L629-R634 delinsW) localized to the dendritic plasma membrane to a lesser extent than wild-type molecules. Quantitative image analysis revealed the differential localization of mutants in PC dendrites (Fig. 8C).

PCs expressing the shRNA-resistant wild-type molecule exhibited normal perpendicular dendrites. In contrast, all disease mutants were defective in the regulation of perpendicular dendrite formation. L253P and E532-M544 del were completely incompetent in perpendicular guidance, while the L629-R634 delinsW mutation, which showed modest mislocalization, retained weak but significant guidance activity (Fig. 8D).

To confirm the effect of the L253P and E532-M544 del mutations in the planar dendrite arborization *in vivo*, we delivered βIII spectrin knockdown plasmid and shRNA resistant βIII spectrin wild-type, L253P or E532-M544 del mutant to immature PCs at E11.5 by *in utero* electroporation. In agreement with *in vitro* observation, L253P and E532-M544 del mutants exhibited abnormal localization in vesicular structures in the soma and proximal dendritic surface, respectively, in contrast to the wild-type molecule spreading over the entire dendritic surface (Fig. 8E). PCs expressing wild-type molecule showed planar dendrites, while the cells expressing either L253P or E532-M544 del mutant displayed disorganized dendrites growing away from the main dendritic plane (Fig. 8E, F). These results suggest that these disease mutations of βIII spectrin disrupt dendritic configuration in PCs.

## Discussion

In the present study, we established a simplified 2D model of axon-dendrite topology using aligned nanofibers and confirmed that PC dendrites grew preferentially in the direction perpendicular to the bundles of afferent parallel fiber axons. The directional arborization is likely a prerequisite for the planar dendrite formation in the cerebellar tissue. Moreover, we revealed that biased dendrite arborization was affected by the loss of αII/βIII spectrins. In control PCs, dendritic branches were formed mainly by terminal bifurcation, with only a few collateral branches emerging from proximal dendrites (Fujishima et al., 2012). In contrast, lateral branching events were significantly increased in βIII spectrin knockdown cells (Fig. 4G).

Dendritic protrusions in differentiating neurons serve as either branch precursors or immature spines (Heiman and Shaham, 2010; Yuste and Bonhoeffer, 2004). Dendrites of developing PCs in culture exhibited numerous lateral protrusions expressing the postsynaptic protein GluD2, suggesting that these lateral protrusions at the proximal dendrites are immature spine precursors. Notably, βIII spectrin knockdown dendrites abnormally extended some proximal protrusions to a length indistinguishable from that of dendritic branches. These elongated protrusions bore multiple GluD2 puncta along their length. Thus, the loss of βIII spectrin seems to alter the fate of proximal dendritic protrusions from immature spines to branch precursors, which become misoriented branches extruded from the main parasagittal plane (Fig. 5).

### Microtubule dynamics and dendritic branching

Branch formation requires microtubule extension into the precursor protrusions or filopodia, which is regulated by dynamic interplays with actin (Burnette et al., 2007; Flynn et al., 2012; Hu et al., 2012). It has been demonstrated that bundled actin in dendritic filopodia guides microtubules into filopodial protrusions from dendritic shafts, while the cortical actin meshwork in the shaft region confines microtubules and suppresses the interaction with bundled actin filaments in filopodia (Dent et al., 2007). We observed frequent microtubule invasion almost exclusively in distal protrusions in normal PCs, supporting the idea that microtubule invasion triggers branch formation and extension predominantly at the distal ends in PC dendrites. βIII spectrin was enriched in the thin neck regions of dendritic protrusions in the dendritic shaft and formed regular or irregular membrane skeletons (Fig. 6H, S7)(Efimova et al., 2017). We assume that βIII spectrin forms the membrane skeletons in coordination with cortical actin that function as a molecular fence dividing dendritic shaft and protrusions, confining microtubule dynamics along dendrites. Thus, in βIII spectrin-deficient PCs, dendritic protrusions with a thin neck were often misshapen with actin signals expanded to the protrusion base and dendritic shaft in contrast to the confined localization in the tip of protrusions in normal cells. Polymerizing microtubules frequently entered these proximal protrusions and promoted their abnormal elongation in βIII spectrin knockdown PCs (Fig. 7).

### SCA5-related mutations affect dendrite growth in PCs

SCA5 is one of the autosomal dominant cerebellar ataxias caused by the heterozygous mutation in βIII spectrin gene (Ikeda et al., 2006). Although our experiment may not completely mimic the disease condition, SCA5 related mutants misexpressed in βIII spectrin knockdown cells failed to substitute for wildtype βIII spectrin to control the dendrite orientation. L253P mutant proteins were not delivered to dendritic membranes nor did they replace the function of wild-type molecules in regulating the oriented growth of PC dendrites. This is consistent with previous studies showing that the L253P mutation affects the trafficking of β-spectrin from the Golgi apparatus (Clarkson et al., 2010). It is also suggested that L253P mutation may reduce the plasticity of actin-spectrin network by enhancing the actin-spectrin affinity. This change might affect the proper dendritic localization of βIII spectrin (Avery et al., 2017). In contrast, the L629-R634 delinsW mutation had only minor effects on dendritic localization and the oriented arborization of dendrites. Interestingly, the E532-M544 del mutant was severely defective in controlling dendrite growth orientation despite its relatively normal dendritic localization except for the distalmost region. It has been proposed that the deletion of E532-M544 possibly affects the triple alpha-helical structures of the spectrin repeats, which might result in the alteration of overall alpha/beta structures (Ikeda et al., 2006). We assume that the E532-M544 del mutation may affect the stabilization of the spectrin architecture in dendrites and interfere with dendrite growth in the normal direction.

The SCA5 patients and βIII spectrin knockout animals exhibit progressive neurodegeneration that has been attributed to the excitotoxicity due to mislocalization and the decreased level of glutamate transporters (Gao et al., 2011; Perkins et al., 2010; Perkins et al., 2016b). Since SCA5 is a late-onset cerebellar ataxia, it may be difficult to directly relate the developmentally disorganized dendrites observed in the present study with symptoms of the adult-onset disease. However, it is suggested that the smaller diameter of dendritic branches in the knockout PCs might alter signal propagation and contribute to the hyperexcitability (Gao et al., 2011). Abnormal growth of dendritic branches in the coronal direction may lead to redundant and inefficient connectivity with parallel fibers (Cuntz, 2012). Furthermore, disruption of planar dendrites is thought to affect the compartmentalization of the cerebellar circuits. In the mature cerebellar cortex, each PC is innervated by a single climbing fiber axon of an inferior olive neuron. Coronally extended PC dendrites invaded in the territory of neighboring PCs might receive abnormal innervation by climbing fibers of neighboring PCs (Gao et al., 2011; Kaneko et al., 2011; Miyazaki and Watanabe, 2011; Perkins et al., 2016a). Developmental defects in the dendrite arborization due to the loss of βIII spectrin might deteriorate the cerebellar function prior to the neuronal degeneration.

### Remaining questions in the perpendicular axon-dendrite interactions

We demonstrate that PC dendrites grow perpendicular to parallel fibers, which seemingly contribute to the planar dendrite arborization *in vivo.* We further present that βIII spectrin is required for the perpendicular dendrite arborization by suppressing the ectopic branch formation in an incorrect direction. However, considering that main dendritic frameworks still form a planar pattern in the βIII spectrin deficient PCs (Fig. 5B), βIII spectrin may function as a gatekeeper to maintain the perpendicular and planar dendrites but not as the main determinant regulating the directional dendrite extension.

The perpendicular interactions may serve as a permissive mechanism for the planar arborization of PC dendrites in parasagittal planes. However, other mechanisms should also be involved in the spatial organization of PC dendrites, as perpendicular contacts would not assure arborization in the exact parasagittal plane. For instance, contact-based repulsion may support the formation of flat dendrites by regulating the space between neighboring dendritic planes (Fujishima et al., 2012; Ing-Esteves et al., 2018) (Fig. 5A).

Nagata et al. (2006) previously proposed that the perpendicular interaction between PC dendrites and parallel fibers might be regulated by the “contact guidance”, where cells recognize the anisotropy of substrates to determine their directionality of the movement. Previous studies using carcinoma cells has demonstrated that anisotropic substrates affect the orientation of focal adhesions and actin fibers anchored to focal adhesions, and thereby lead to the directional cell movement (Ray et al., 2017). We assume that PC dendrites might sense the anisotropy of the GC axons and adjust the orientation of the adhesions and cytoskeletal components including spectrin tetramers, thereby affecting the growth orientation of dendrites.

Previous studies using diffusion magnetic resonance imaging have revealed a 3D grid-like organization of axonal bundles in the forebrain, in which two distinct fibers interweave and cross at nearly right angles (Wedeen et al., 2012). Such a grid-like organization might be formed by perpendicular contact guidance between different types of axons. It is of interest to confirm whether perpendicular contact is a general mechanism in neural network formation.

## Materials and methods

### Antibody list

|  |  |  |
| --- | --- | --- |
| Rabbit polyclonal anti-βIII Spectrin | SantaCruz | sc-28273 |
| Goat polyclonal anti-βIII Spectrin | SantaCruz | sc-9660 |
| Mouse monoclonal anti-βIII Spectrin | SantaCruz | sc-515737 |
| Mouse monoclonal anti-αII Spectrin | SantaCruz | sc-376849 |
| Rabbit polyclonal anti-GFP | Thermo Fisher Scientific | A-11122 |
| Mouse monoclonal anti-GFP | Thermo Fisher Scientific | A-11120 |
| Rabbit polyclonal anti-RFP | MBL Life Science | PM005 |
| Mouse monoclonal anti-Calbindin | Swant | 300 |
| Mouse monoclonal anti-c-Myc | SantaCruz | sc-40 |
| Mouse monoclonal anti-HA | Roche | 1583816 |
| Rabbit polyclonal-Pax6 | Wako | 019-27291 |
| Rabbit polyclonal anti-GluD2-C | Frontier Institute | GluRD2C-Rb-Af500 |
| Mouse monoclonal anti-tau | MERCK | MAB3420 |
| Donkey anti-Mouse Alexa Fluor 488 | Thermo Fisher Scientific | A-21202 |
| Donkey anti-Mouse Alexa Fluor 555 | Thermo Fisher Scientific | A-31570 |
| Donkey anti-Mouse Alexa Fluor 647 | Thermo Fisher Scientific | A-31571 |
| Donkey anti-Rabbit Alexa Fluor 488 | Thermo Fisher Scientific | A-21206 |
| Donkey anti-Rabbit Alexa Fluor 555 | Thermo Fisher Scientific | A-31572 |
| Donkey anti-Rabbit Alexa Fluor 647 | Thermo Fisher Scientific | A-31573 |
| Donkey anti-Goat Alexa Fluor 555 | Thermo Fisher Scientific | A-21432 |
| Donkey anti-Goat Alexa Fluor 568 | Thermo Fisher Scientific | A-11057 |
| Donkey anti-Goat Alexa Fluor 647 | Thermo Fisher Scientific | A-21447 |

### Mice

Mice were handled in accordance with the guidelines of the Animal Experiment Committee of Kyoto University and were housed in a dedicated pathogen-free environment with a 12-hour light/dark cycle.

### Plasmids

The pAAV-CAG-GFP (or mCherry)-hH1 vector, including the human H1 promoter, was used to express shRNA to knockdown target gene expression as previously described (Fukumitsu et al., 2015). The targeting sequences were designed by using the web-based software siDirect (Naito et al., 2009): control shRNA (5’-GCATCTCCATTAGCGAACATT-3’), βIII-spectrin shRNA (5’-GTCAATGTGCACAACTTTACC-3’), αII spectrin shRNA (5’-GTAAAGACCTCACTAATGTCC-3’). To generate resistant mutants of αII spectrin and βIII-spectrin that contained three silent mutations within shRNA target sequences, the cDNA of mouse αII spectrin or βIII-spectrin was cloned from a mouse brain cDNA library and mutagenized by using a PCR-based method. To generate the L253P, E532-M544 del and L629-R634 delinsW βIII spectrin mutants, PCR-based mutagenesis was performed by using the resistant mutant of βIII-spectrin as a template. αII spectrin tagged with HA at the N-terminus and βIII-spectrin wild-type and mutant sequences tagged with myc at the N-terminus were cloned into the pCAGGS vector. To generate the EB3-EGFP construct, the coding sequence of EB3 was amplified from a mouse brain cDNA library and inserted into the pAAV-CAG-EGFP plasmid. For the CRISPR/Cas9-based knockout of βIII-spectrin, the guide RNA sequence was selected by using the web-based software CRISPRdirect (Naito et al., 2015). The βIII-spectrin target sequence (5’-GAGACCTGTACAGCGACCTG-3’) was inserted into pSpCas9(BB)-2A-GFP (PX458) (Addgene plasmid #48138) (Ran et al., 2013).

### *In utero* electroporation

The *in utero* electroporation of plasmids was performed as described previously (Nishiyama et al., 2012). Briefly, pregnant mice on day 11.5 of gestation were deeply anesthetized via the intraabdominal injection of a mixture of medetomidine, midazolam, and butorphanol. Plasmid DNA (1-5 μg/μl) was microinjected into the fourth ventricle of the embryos (FemtoJet; Eppendorf). Then, five current pulses (amplitude, 33 V; duration, 30 ms; intervals, 970 ms) were delivered with a forceps-shaped electrode (CUY650P3; NepaGene) connected to an electroporator (CUY21; NepaGene).

### *In vivo* electroporation to label parallel fibers

The *in vivo* electroporation of plasmids was performed as described previously (Umeshima et al., 2007). P8 ICR mice were cryoanesthetized. A small burr hole was made in the skull over the cerebellum with a 27-gauge needle. The plasmid DNA (pAAV-CAG-mCherry) was microinjected through the hole by using a syringe with a 33-gauge needle (Ito). A forceps-shaped electrode connected to the cathode of an electroporator (CUY21; NepaGene) was placed in the occipital region. A needle used for DNA injection was connected to the anode. Then, six current pulses (amplitude, 70 V; duration, 50 ms; intervals, 150 ms) were delivered. After the wound was sutured, the pups were warmed at 37°C and returned to the home cage.

### Primary cerebellar culture and nucleofection of cerebellar neurons

The primary culture of cerebellar neurons was performed as previously described (Fujishima et al., 2012) with slight modifications. Cerebella from postnatal day (P) 0 mice were dissected in HBSS (GIBCO) and dissociated using a Neuron Dissociation Kit (FUJIFILM Wako Pure Chemical Corporation). Cells were plated on a 12 mm coverslip coated with poly-D-lysine in DMEM/F12 supplemented with 10% FBS at a density of 1.5 cerebella/coverslip. Following incubation, the media were replaced with maintenance media containing DMEM/F12, 0.1 mg/ml bovine serum albumin, 2.1 mg/ml glucose, 2X Glutamax, 8 μM progesterone, 20 μg/ml insulin, 200 μg/ml transferrin, 100 μM putrescine, 30 nM selenium dioxide, 4 μM AraC and 1% penicillin-streptomycin. For cultures on nanofibers, dissociated neurons were plated on aligned or random nanofiber plates (Nanofiber solutions).

For the transfection of plasmid DNA into PCs, nucleofection was performed as described previously (Kawabata Galbraith et al., 2018). Briefly, dissociated cerebellar cells from 1.5-2 cerebella were washed twice with OptiMEM and resuspended in 100 μl of OptiMEM containing 5-8 μg of plasmid DNA. Cells were transferred to a cuvette and nucleofected with a Nepa21 electroporator (NEPAGENE).

To separate large cell fraction (containing PCs, interneurons and glia) and small cell fraction (containing GCs), dissociate cells from P0 cerebellar tissue were purified with a step gradient of Percoll (Hatten, 1985). Isolated GCs from P0 mice were plated on the aligned fiber in a density of 10 x 10^5^, 7 x 10^5,^ and 5 x 10^5^cells/cm^2^. Then, a large cell fraction including PCs was plated on the culture with a density of 10 x 10^4^, 7 x 10^4,^ and 5 x 10^4^cells/cm^2^, respectively.

For the coculture of cerebellar microexplant and PCs, a microexplant culture from cerebellar tissues was prepared as described previously (Nakatsuji and Nagata, 1989). Briefly, the external granular layer of the cerebellar cortex from P2 mice was dissected into 300-500 μm pieces and plated on the poly-d-lysin and laminin-coated coverslips. One day after plating, the isolated large cell fraction including PCs from P0 mice was added to the culture.

### Immunofluorescence and image acquisition

For immunocytochemistry, cells cultured on coverslips or nanofibers were fixed for 15 min at RT in 4% paraformaldehyde (PFA) in PBS. Cells were washed and permeabilized with PBS containing 0.25% Triton (PBS-0.25T). The cells were then blocked with a blocking solution (PBS (−0.25 T) with 2% bovine serum albumin (BSA)) for 30 min at RT. The cells were incubated with primary antibodies at 4°C overnight in blocking solution, washed with PBS, and incubated with fluorescently labeled secondary antibodies in blocking solution.

For immunohistochemistry, the mice were anesthetized with isoflurane and perfused with PBS followed by 4% PFA in phosphate buffer. Their brains were removed and postfixed overnight in 4% PFA/PBS at 4°C. After washing with PBS, the brains were embedded in a 3.5% low-melt agarose in PBS. Sagittal or coronal sections were cut at a 50-100 μm thickness by using a vibratome (Dosaka). The sections were permeabilized in PBS with 0.5% Triton (PBS-0.5 T) and blocked with 2% skim milk in PBS (−0.5 T) for 30 min. The sections were incubated with primary antibodies at 4°C overnight in blocking solution, washed with PBS, and incubated with fluorescently labeled secondary antibodies in 2% skim milk in PBS (−0.5 T).

Images of the fixed samples were acquired on a laser scanning confocal microscopy Fluoview FV1000 (Olympus) equipped with UPLSAPO 40x dry (NA 0.95), 60x water-immersion (NA 1.2) and 100x oil-immersion objectives (NA 1.40). For the imaging of PC dendrites in the cerebellar tissue, serial confocal z stack images were acquired from midsagittal regions in lobes IV-V with the 60x objective at a z-step of 0.57 μm. The images were three-dimensionally reconstructed by using Imaris software (Bitplane). For the imaging of PC dendrites in the dissociated culture, serial confocal z stack images were obtained with 40 x or 100 x objectives.

### Quantification of dendrite morphology

For the analysis of dendritic flatness, captured confocal z-stacked images were binarized and skeletonized in ImageJ *(“Skeletonize (2))/3)))”* plugin). To identify the plane with the closest fit to the given dendritic arbor, a principal component analysis was performed in MATLAB software (Fig. S2). To analyze the distance of the dendritic branches from the fitted plane, 3000 points (for Fig. 1A, B) or 100 points (for Fig. 1D, 3E, 8E) in the skeletonized dendritic images from each cell were randomly selected, and the distance between each point and the fitted plane was calculated.

For the morphometric analysis of PC dendrites on aligned fibers, Z-projected images were binarized and skeletonized in the Image J plugin or MATLAB software. To analyze the branch angle, dendritic branches were divided into 3.5 μm segments, and the angle between each segment and fiber was quantified (Fig. S3).

### STED imaging

We used a Leica TCS SP8 STED with an oil immersion 100x objective lens with NA 1.4 (HC-PL-APO 100x/1.4 OIL, Leica) to analyze the subcellular localization of βIII spectrin. The protein was labeled with anti-βIII spectrin (Santa Cruz, SC-28273) and a secondary antibody conjugated with Alexa555 (Thermo Fisher). The fluorophore was excited with a white laser tuned to 555 nm and depleted with a 660 nm STED laser. A time gate window of 0.35-3.85 ns was used to maximize the STED resolution.

### Time-lapse imaging, image processing, and image analysis

For long-term time-lapse imaging, fluorescently labeled PCs were observed every 1-3 hours with an incubation microscope (LCV100, Olympus) equipped with a 20x objective (NA 0.7, Olympus). Serial z-stacked images were obtained at z-steps of 1 μm (1 μm x 5 steps). For the high-resolution live imaging of dendritic arbors (used for EB3 imaging experiments),

For the analysis of EB3 dynamics, We used 10-11 DIV PC dendrites transfected with pAAV-CA-EB3-EGFP and pAAV-CAG-mCherry/hH1-control or hH1-βIII spectrin. Time-lapse images were obtained with a confocal microscope (FV1000, BX61W1; Olympus) equipped with a LUMFI 60x objective (NA 1.10, Olympus) under 5% CO2 supplementation. Images were recorded with 3x digital zoom at an interval of 3 s. To eliminate background cytosolic signals average subtraction was performed (Schätzle et al., 2016), in which an average projection of all of the time-lapse images of EB3-EGFP signals was generated and subtracted from each frame of the time-lapse images. The images were then processed via an unsharp masking procedure in ImageJ software to obtain the enhanced images. For the quantitative analysis, the number of dendritic protrusions in the dendritic segments of 10 μm at the most distal regions or proximal regions (more than 10 μm away from the most distal end) were counted in imageJ. Of those, dendritic protrusions invaded by the EB3-EGFP were identified manually with the assistance of imageJ plugin TrackMate (Tinevez et al., 2017).

## Acknowledgments

We thank iCeMS Analysis Center and Dr. Yukiko Ohara for technical assistance.

## Author Contributions

Conceptualization, K.F. and M.K.; Methodology, K.F. and M.K.; Investigation, K.F., J.K. and M.Y.; Formal analysis, K.F., J.K. and M.Y.; Writing K.F. and M.K.

## Supplementary Figure legends

Fig. **S1 Validation of shRNA targeting βIII spectrin in dissociation culture**

Representative images of cerebellar PCs transfected with a plasmid containing EGFP/shRNA control (ctr) or EGFP/shRNA βIII spectrin (βIII kd). Cells were stained with anti-calbindin (magenta) and anti-βIII spectrin (white) antibodies. The calbindin-positive PCs transfected with the βIII knockdown plasmid lacked βIII spectrin signals. Scale bars: 20 μm.

**Fig. S2 Quantitative analysis of the planarity of PC dendrites in cerebellar tissue**

**(A)** Representative image of cerebellar PCs labeled with GFP in a sagittal cerebellar slice at P14. **(B)** Skeletonized image of the PC dendrites in (A). Magenta points on the skeletonized dendrite were randomly selected. **(C)** A three-dimensional view of the skeletonized dendrite, randomly selected points, and a plane fitted to the dendrites. **(D)** A side view of randomly selected points and the dendritic plane.

**Fig. S3 Quantitative analysis of dendrite orientation in the culture using nanofiber substrates**

(A) Representative image of PCs growing on the aligned nanofiber substrates. (B) A DIC image showing nanofibers. (C) Traced dendrites were divided into 3.5-micron segments. The angles of dendritic segments with respect to the average orientation of nanofibers were recorded. (D) The traced dendrites were pseudocolored by the orientation of the dendritic segments.

**Fig. S4 Purkinje cell dendrites grow perpendicular to granule cell axons but not nanofibers**

(A) Representative images of PCs (calbindin) grown on microexplant culture. Asterisks indicate the locations of microexplants. GCs extend their axons radially from the explants (tau). One day after the preparation of microexplants, large cells including PCs were isolated from cerebellar tissue by the Percoll gradient methods and plated on the explant. Microexplants and PCs were fixed at 9 days after the preparation of microexplants. Right panels were magnified views of the boxed region in the right image. (B) Surface rendering images showing the PC and GC axons in (A, middle).

**Fig. S5 The growth of Purkinje cell dendrites on the axonal fibers with different densities**

(A) Representative images of PCs (calbindin) and GC axons (tau) grown on nanofiber substrates. GCs isolated by the Percoll gradient method were plated on the aligned nanofiber substrates at the density of 10 x 10^5^/cm^2^, 7 x 10^5^/cm^2^, or 5 x 10^5^/cm^2^. Large cells including PCs were plated at the density of 10 x 10^4^/cm^2^, 7 x 10^4^/cm^2^, or 5 x 10^4^/cm^2^, respectively. (B) Magnified views of dendrites in the boxed regions in (A). Note that PCs cannot extend their dendrite in the region where the density of GC axons is low, even though the region contains a comparable level of nanofiber substrates (asterisk). Scale bars: 20 μm.

**Fig. S6 CRISPR/Cas9-based βIII spectrin knockout affects the perpendicular interaction between PC dendrites and GC axons.**

Representative images of PCs on aligned fibers transfected with CRISPR/Cas9 vectors (left: control empty vector, right: βIII spectrin target sequence). Cells were stained with anti-βIII spectrin (magenta). The βIII spectrin knockout cell exhibits a dendrite elongating in an abnormal orientation. n = 22 cells for control, 20 cells for βIII spectrin knockout. Scale bar: 30 μm.

**Fig. S7 Membrane periodic structure of βIII spectrin in PC dendrites**

**(A and B)** Representative STED image of βIII spectrin in the proximal region (A) and distal region (B) of developing PC dendrites at 9-11DIV. **(C)** ROI1-3 (ROI1: dendritic shaft in (A), ROI2-3: dendritic filopodia in (B)) are magnified views. The normalized intensities of βIII spectrin repeats in ROIs 1-3 are shown by line plots below. Gray dotted lines are drawn every 185 nm from the first peak of the βIII spectrin signal. **(D)** The histogram indicates the distribution of the spacing of repeated βIII spectrin structures in the dendritic shaft (blue) and the base of dendritic protrusions (red). Shaft: n = 85 from 11 dendrites. Base of protrusions: n = 103 from 13 dendrites (dendritic base). **(E)** Another example of a STED image of βIII spectrin in the shaft region of a developing PC dendrite. **(F)** ROI4-6 are magnified views of the boxed regions in (A). Normalized intensities of βIII spectrin repeats are shown by line plots below. Scale bars: 2 μm in (A), (B) and (E). 200 nm in (C) and (F).

