## Supplementary figures and images for "βIII spectrin controls the planarity of Purkinje cell dendrites by modulating perpendicular axon-dendrite interaction"

### Supplemental Figures

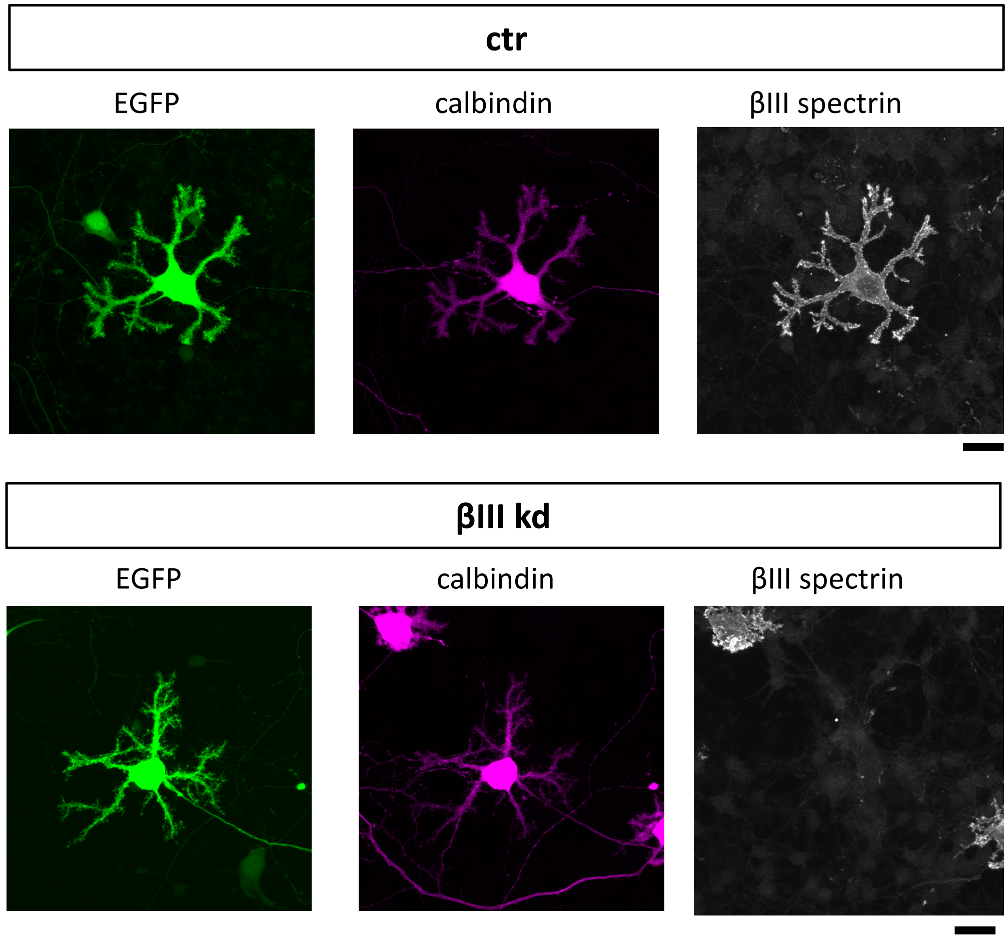

#### Figure S2

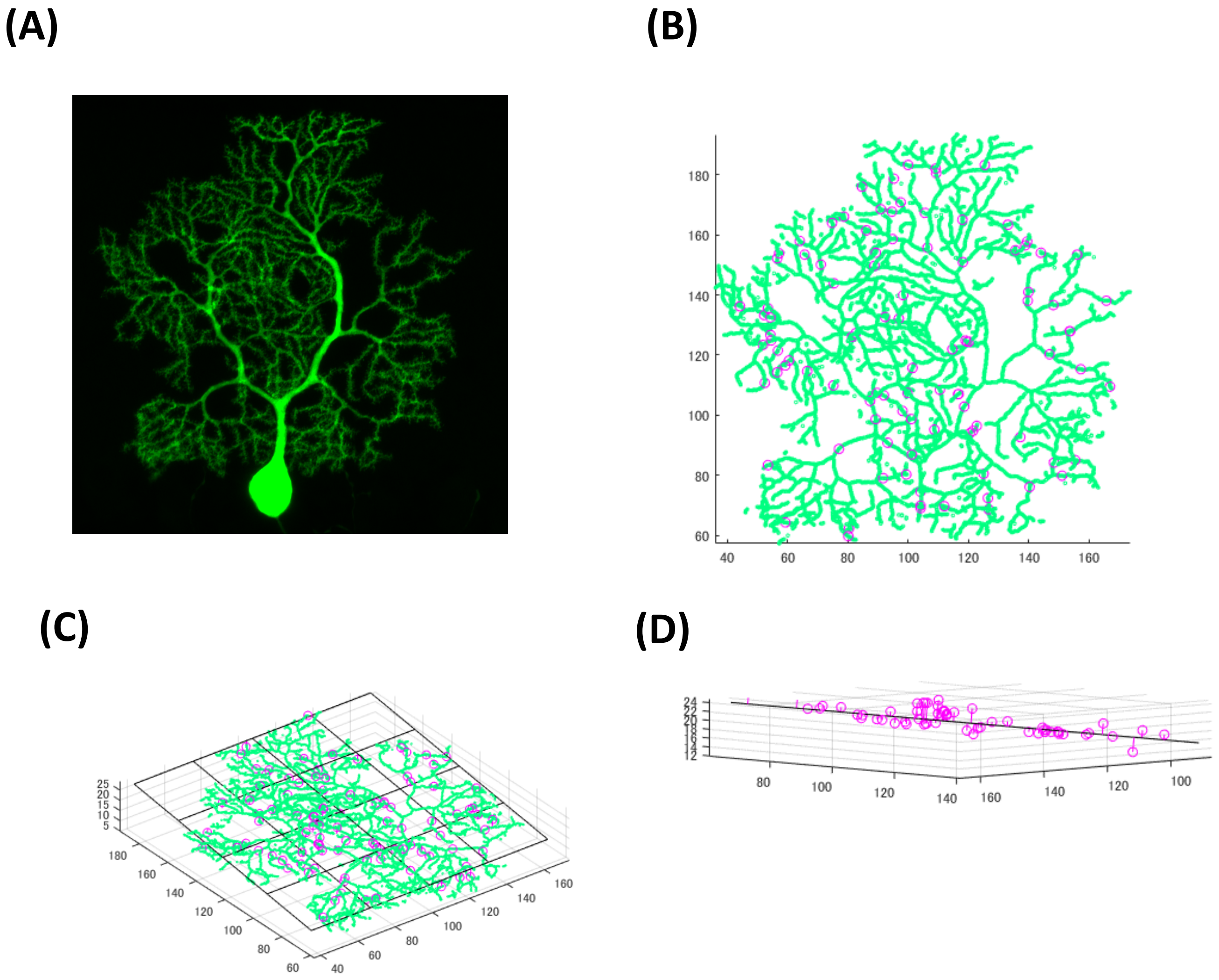

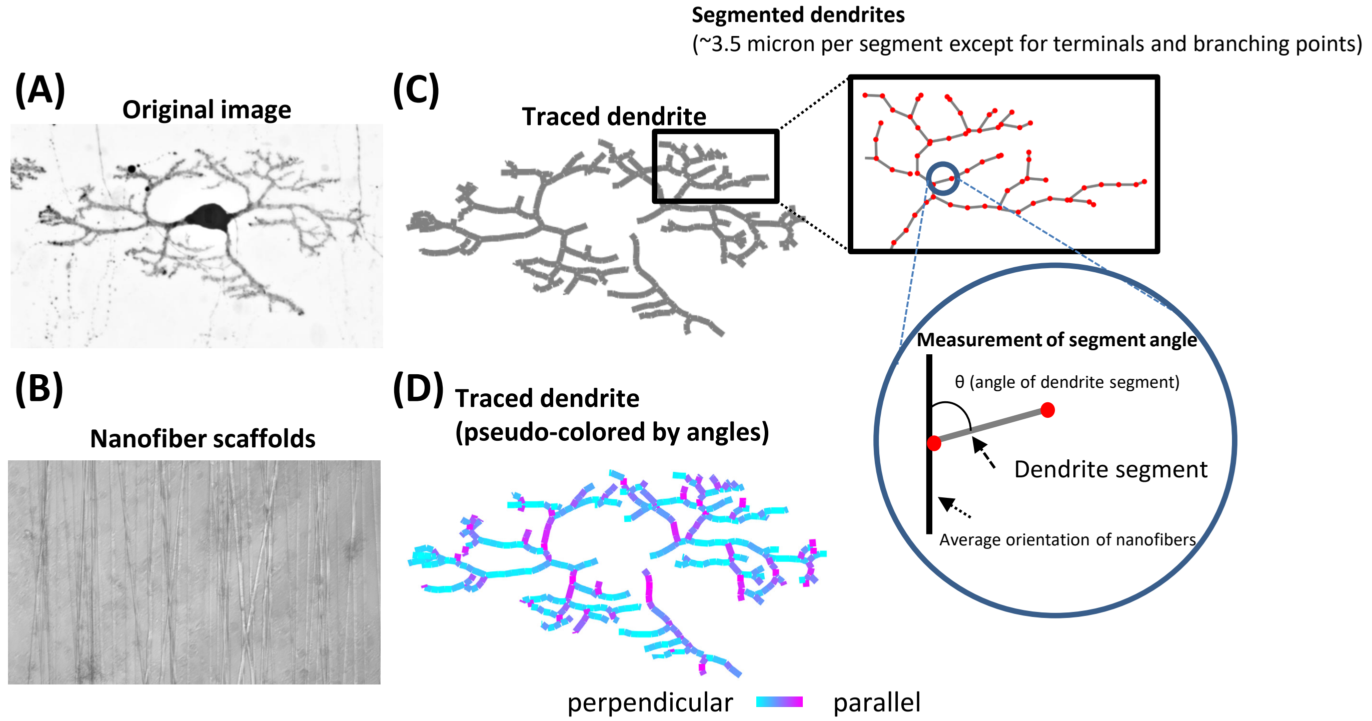

#### Figure S4

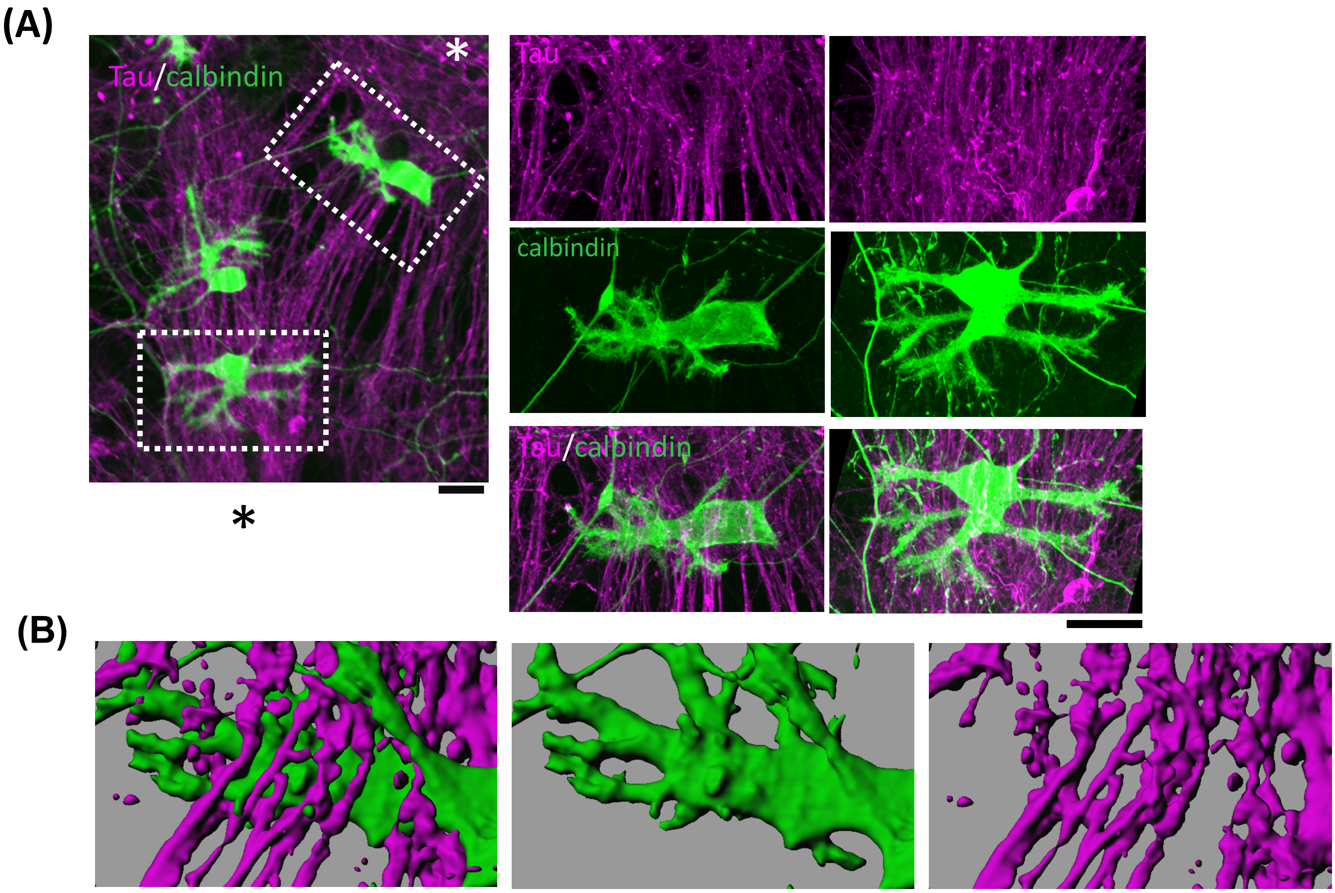

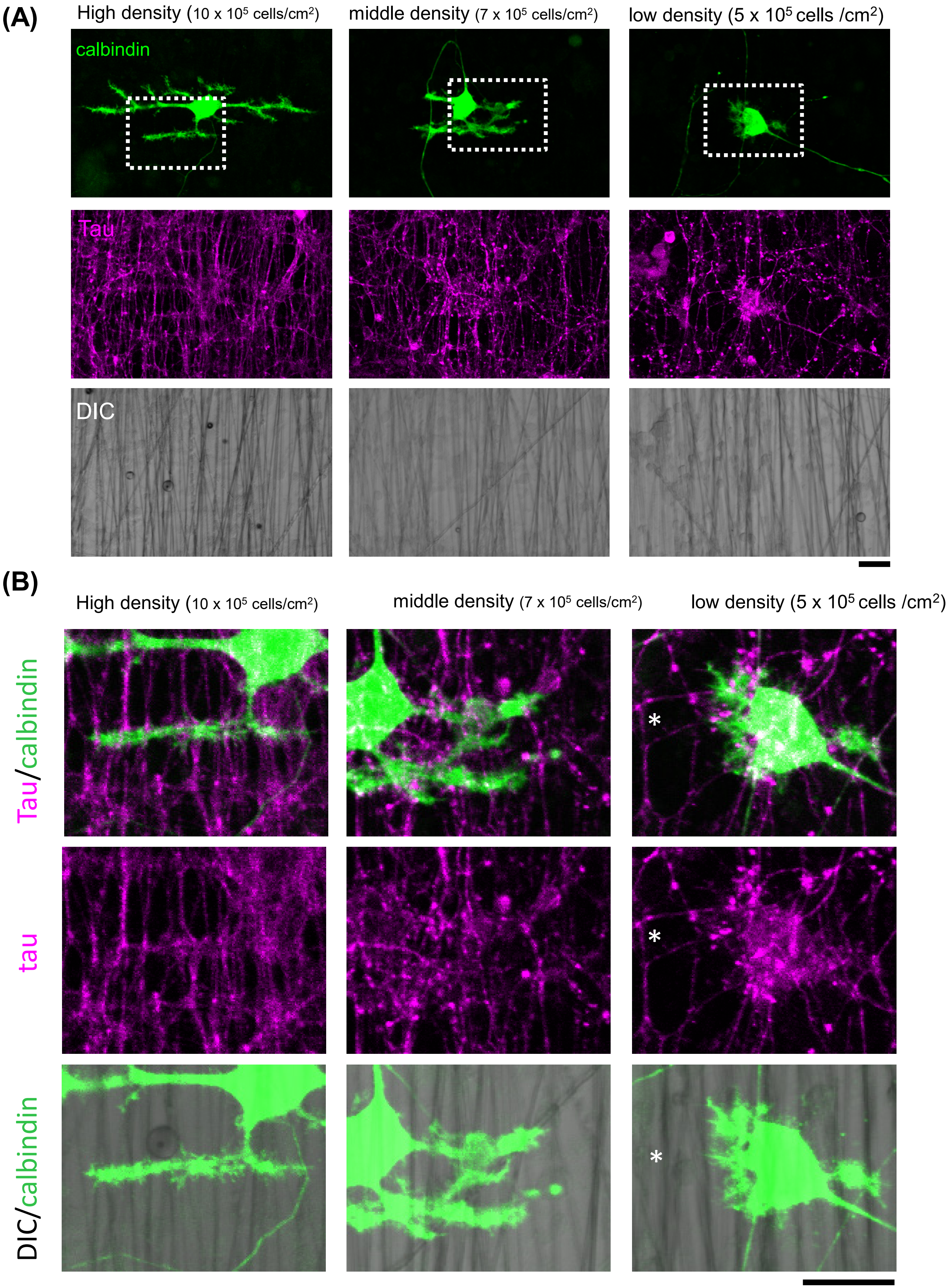

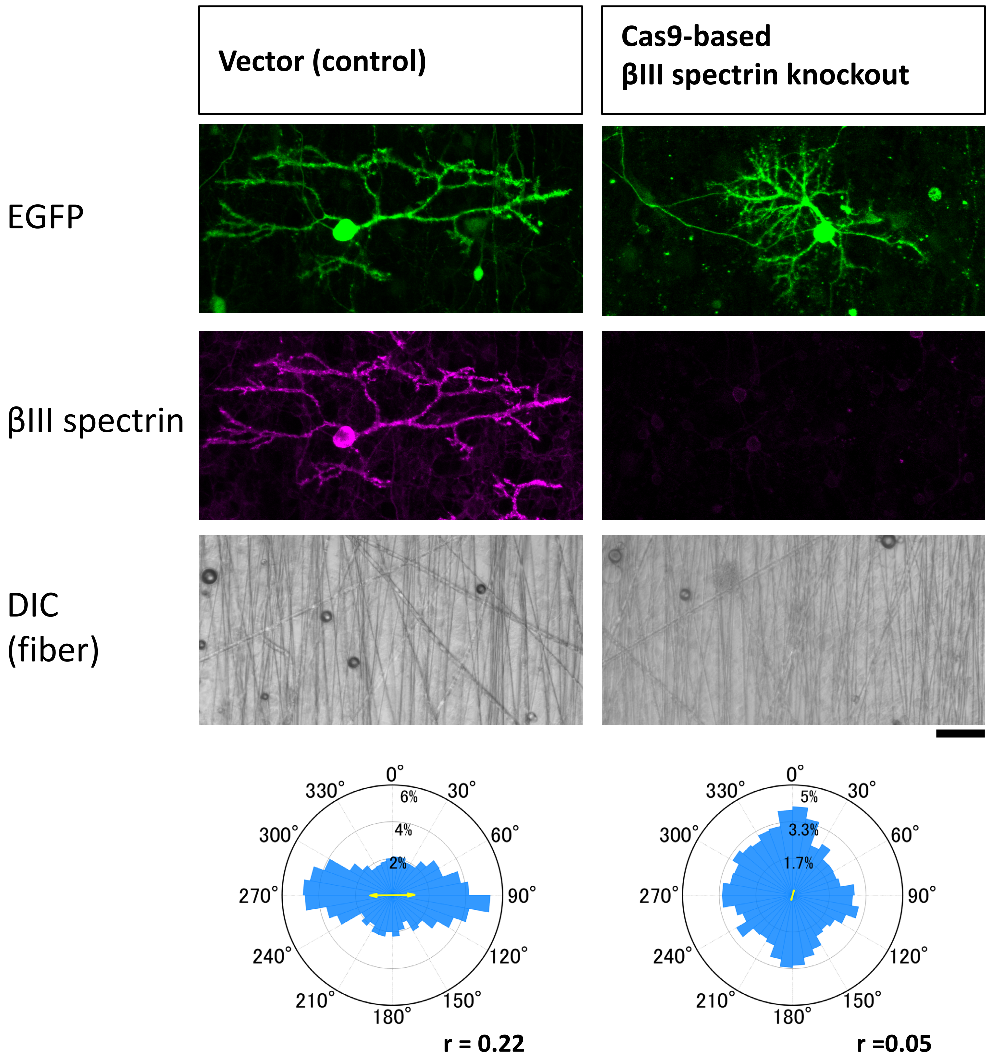

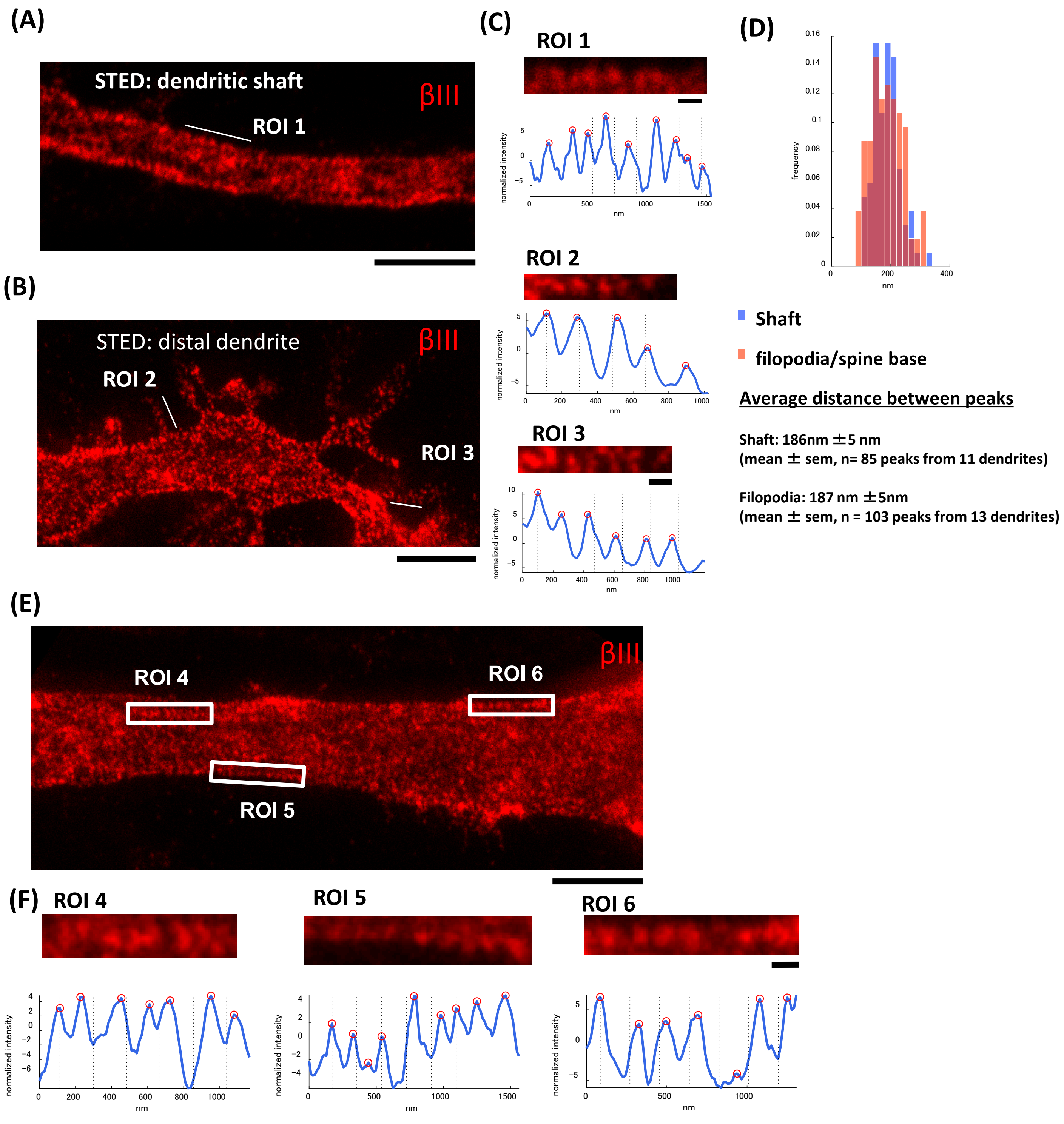
